# Phosphoproteomic Heterogeneity Reveals p-SRSF3-Driven Metastasis in Early Hepatocellular Carcinoma

**DOI:** 10.1101/2025.02.11.637605

**Authors:** Liangliang Ren, Jiahan Jiang, Changwen Ning, Weiran Lin, Lihui Qi, Chaoying Li, Ying Jiang

## Abstract

Mechanisms driving progression and metastasis in early-stage hepatocellular carcinoma (HCC) remain poorly clarified. Although phosphoproteomics has been applied to broader HCC cohorts, the heterogeneity of phosphorylation signaling in early-stage HCC remains unexplored. Here, we conducted integrated phosphoproteomic and proteomic analysis on early-stage HCC tumors and paired non-tumor tissues. By comparing tumor and non-tumor tissues, we identified markedly reduced phosphorylation of EGFR-mediated canonical ERK/MAPK signaling, alongside elevated atypical MAPK signaling in HCC. To further explore the signaling heterogeneity within tumors, we applied nonnegative matrix factorization (NMF) and consensus-clustering, and identified three distinct phosphoproteomic subtypes with unique phosphorylation signatures and clinical characteristics. Interrogating the most malignant subtype, we found markedly high abundance of phosphorylation of mRNA splicing regulators, and an enrichment of epithelial-mesenchymal transition (EMT) pathways in the proteome, with phosphorylation of the splicing factor SRSF3 emerging as a critical feature. Mechanistically, we show that SRSF3 hyperphosphorylation, driven by SRPK1/CLKs and potential PP1 phosphatase suppression, promotes metastasis and proliferation. Critically, SRPK1/CLKs inhibitors effectively suppressed SRSF3 hyperphosphorylation-mediated migration and invasion of HCC cells. Our study uncovers a new dimension of signaling-driven tumor heterogeneity in early-stage HCC and identifies the SRPK1/CLKs-SRSF3 hyperphosphorylation axis as a key metastatic driver, suggesting a potential therapeutic target for early intervention in HCC.

## Introduction

HCC is the third deadliest cancer worldwide, with a 5-year survival rate of 18% [1]. Metastases rather than the primary tumors are an overwhelming major contributor to canerassociated deaths [2]. The high heterogeneity in HCC limits effective therapies [3], particularly in early stages, where the mechanisms of initiation, progression, and metastasis remain unclear. Protein phosphorylation, coordinated by kinases and phosphatases, dramatically expands the diversity of proteins and regulates signaling. Previous studies have focused on the phosphorylation signatures that vary in distinct proteomic subtypes [4–6] or signaling pathways at the collapsed protein phosphorylation level and revealed subgroups consistent with the proteomic subgroups [7]. However, the phosphoproteome in early-stage HCC has not been extensively explored. Interrogation of the phosphoproteome in early-stage HCC provides a valuable window into the oncogenic signaling events that may drive tumor initiation, early progression, and metastasis, which can potentially uncover the phosphoproteomic signatures that resolve the heterogeneity of patients with early-stage disease, or signaling pathways and targets for interventions that prevent progression to more aggressive HCC.

In this study, our integrated phosphoproteomic analysis on early-stage HCC identifies dysregulated phosphorylation signatures, and stratifies the patients based on tumor heterogeneity. The phosphoproteomic subtypes identified molecular signatures of malignant transformation and mechanisms contributing to HCC progression and metastasis in the early stage, which should, in turn, expedite clinical classification and targeted therapies for patients with early-stage HCC.

## Results

### Phosphoproteomic characterization of early-stage HCC

In 103 paired tumor and adjacent non-tumor tissues of clinical early-stage hepatocellular carcinoma related to hepatitis B virus infection, phosphoproteomics identified 27,651 class I phosphosites (localization score *>*0.75) and 9,254 proteins [4] Quanlity control was performed using ROBPCA [8], and six paired samples were identified as outliers and excluded from further analysis (Figure S1A). Among the remaining 97 pairs of samples, 88 had the corresponding proteomic expression data (Table S0). Principal component analysis revealed a clear separation between tumor and non-tumor samples at the phosphoproteome level, while also highlighting the heterogeneity in tumor samples (**Figure 1**A). We then examined the correlation between the abundance of 27,651 phosphosites and their aggregated protein phosphorylation levels with the corresponding protein abundance in the 88 paired tumor and non-tumor samples, respectively. In non-tumor tissue samples, 7,593 of the 27,651 phosphosites and their corresponding protein pairs were analyzed (Figure 1B). The overall correlation between the phosphoproteomic and proteomic data was relatively weak, with a median Spearman’s r value of 0.076; only 1.6% showed a significant positive correlation (Benjamin-Hochberg *adj.p*-value *<*0.05). In the tumor tissues, 8,912 of the 27,651 phosphosites and their corresponding protein pairs were analyzed, presenting a median Spearman’s r value of 0.169 of the overall correlations, and 9.7% showed a significant positive correlation (Benjamin-Hochberg *adj.p*-value *<*0.05). Even when considering aggregated protein phosphorylation levels from these phosphosites, only 6.7% and 24.5% of the correlations showed significant positive correlation in non-tumor and tumor tissues, respectively (Figure S1B). Gene set enrichment analysis (GSEA) revealed that proteins with high correlations between protein expression and phosphorylation were enriched in hallmarks of glycolysis and EMT, whereas those with low correlations were enriched in cellular signaling transduction, such as signaling by GPCR(Figure 1C, Table S1). Overall, these distinct correlation patterns across functional pathways likely reflect a broad post-translational regulatory shift of cellular signaling in HCC.

**Figure 1:**
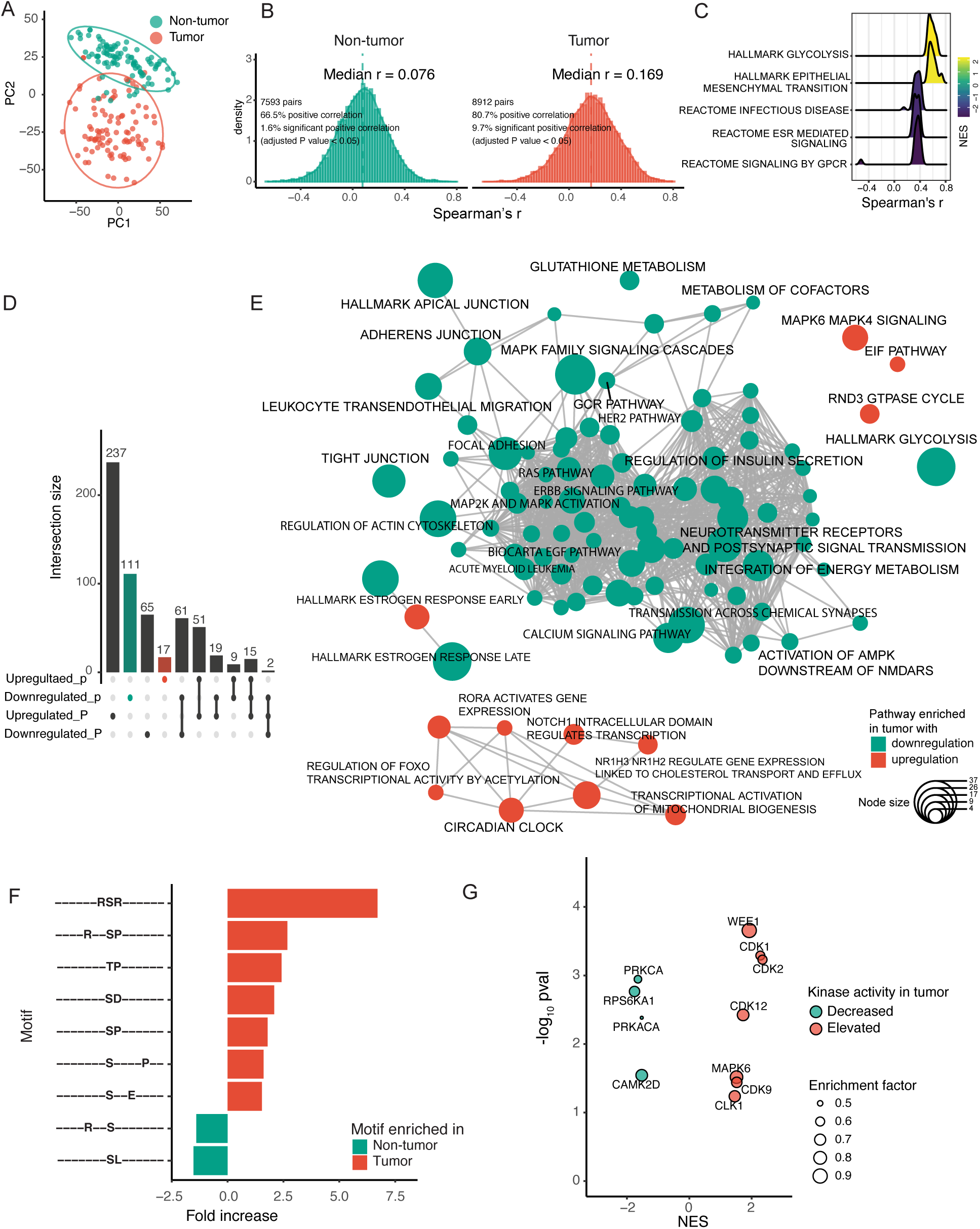
(previous page.) The dysregulated protein phosphorylation in early-stage HBV-related HCC cohort (Jiang et al.’s cohort) (A) Principal component analysis of the phosphoproteome in the tumor and non-tumor tissues. (B) Correlation of levels between phosphorylation at phosphosite and protein expression in the non-tumor and tumor tissues (Benjamin-Hochberg adjusted *p*-value *<*0.05). (C) GSEA enrichment analysis of pathways and hallmarks for proteins with high or low correlations of protein expression and phosphorylation level in tumor tissues. (D) UpSet plot showing the number of pathways and hallmarks enriched from regulated proteins and phosphoproteins. Pathways and hallmarks that were upregulated or downregulated in the phosphoproteome are denoted as “Upregulated p” or “Downregulated p”, respectively, while those in the proteome are denoted as “Upregulated P” or “Downregulated P”. Pathways and hallmarks observed in one or multiple conditions are connected by black dots. The bar plot of pathways and hallmarks only observed with downregulation of phosphoproteins is highlighted with dark green, while that only observed with upregulation is highlighted with dark red. (E) Visualization of pathways and hallmarks that were significantly enriched and only observed with dysregulated protein phosphorylation in the tumor tissues. The similarity of the pathway enrichment is calculated by the Jaccard correlation coefficient method, and the node size indicates the number of overlapping proteins between gene sets. (F) Bar plot showing motif fold changes for the upregulated and downregulated phosphosites (compared to all observed phosphosites). The upregulated phosphosites were enriched with basophilic motifs (XRSRX, RXXSP), proline-directed motifs (XXTP and XXSP), and acidophilic motifs (SXXE, XXSD). Those downregulated were enriched with motifs of RXXS and XXSL. (G) Kinase-substrate enrichment analysis (KSEA) showed decreased and elevated kinase activities in tumor, with corresponding normalized enrichment score (NES). The size of the kinase indicates the enrichment factor.

### The dysregulated phosphorylation signatures and signaling pathways involved in carcinogenesis and progression of early-stage HCC

Differentially expressed phosphorylation site (DES) analysis of the phosphoproteome revealed 1,950 upregulated and 1,660 downregulated phosphosites (Table S2), and differentially expressed protein (DEP) analysis of the proteome showed 2,949 upregulated and 679 downregulated proteins (Table S5) in early-stage HCC. Enrichment analysis of biological processes for proteins with DES revealed that mRNA processing, chromatin remodeling, and RNA catabolic processes are regulated by elevated phosphorylation in HCC, whereas metabolic processes are regulated by decreased phosphorylation (Figure S1C, Table S3). In terms of signaling pathways and hallmarks, mRNA splicing, spliceosome, Rho GTPase cycle, RNA polymerase II-mediated transcription, and transport of mature transcript to cytoplasm , etc., were regulated by elevated phosphorylation in HCC (Figure S1D, Table S4). Correspondingly, the insulin signaling pathway and metabolic pathways such as metabolism of glucose, carbohydrates, amino acids, and derivatives showed decreased phosphorylation regulation in HCC (Figure S1D, Table S4), indicating reduced physiological functions related to liver metabolism.

By comparing the signaling pathways and hallmarks that were significantly regulated at the protein expression and phosphorylation level in early-stage HCC (Figure 1D, see also Table S6, S7 and S8), 51 hallmarks and pathways were observed with both elevated protein expression and phosphorylation, including the pathways involved in chromatin modifying enzymes, mRNA processing, cell cycle, E2F and MYC targets (Table S7). Meanwhile, 61 hallmarks and pathways were observed with both decreased protein expression and phosphorylation (Figure 1D, see also Table S7), including pathways involving lipid metabolism such as fatty acid metabolism, bile acids and bile salts, metabolism of carbohydrate, xenobiotics, and amino acid and derivatives, highlighting the loss of metabolism functions of liver in HCC. In addition, 17 and 111 pathways or hallmarks were observed to be solely regulated by elevation or reduction at the phosphorylation level (Figure 1D). To identify functional modules of pathways that were only observed in the phosphoproteome, we created an enrichment map based on the UpSet plot of pathway enrichment results (Figure 1E, Table S8). One cluster and several pathways were identified in the pathways with elevation only on phosphorylation regulation, these include the circadian clock, transcriptional activation of mitochondrial biogenesis, and gene expression regulated by RORA, FOXO, NR1H3/NR1H2, EIF pathway, RND3 (RHOE) GTPase cycle, MAPK6 and MAPK4 signaling, etc. (Figure 1E) . In contrast, a network with two clusters involved in metabolic and signaling pathways was observed with only a decrease in phosphorylation regulation, one of which consisted of metabolism of cofactors, AMPK downstream signaling, neurotransmitter receptors, calcium signaling pathway, and regulation of insulin secretion. Another cluster consisted of modules on cell adhesion, migration, and cancer-related signaling pathways, including hallmark apical junction, adherens junction, regulation of actin cytoskeleton, signaling of MAP2K, MAPK family cascades, ERBB, HER2, EGF, and hallmark of the estrogen response (Figure 1E, Table S8). Overall, the dysregulated hallmarks and pathways indicated post-translational regulation by phosphorylation in HCC.

To further explore the kinases that may determine the dysregulated signaling in HCC, by analyzing with the Motif-X algorithm on the amino acid sequences surrounding the 1,950 elevated phosphorylation sites and the 1,660 decreased phosphorylation sites [9], several basophilic kinase motifs, such as XRSRX (SRPKs [10]), RXXSP (SRPKs, CLKs [10]), and proline-directed motifs XXTP and XXSP (MAPK family kinases [11] and CDK family kinases [11]), as well as acidophilic kinase motifs, such as SXXE and XXSD (CK1 [11], CK2 [11, 12]) were significantly enriched in the elevated phosphorylation sites in early-stage HCC (Figure 1F, Table S9), suggesting that the activities of SRPKs and CLKs, members of MAPKs, CDKs, CK1, and CK2 family kinases were likely elevated in HCC. In contrast, phosphosites with decreased levels in HCC were enriched in motifs of RXXS (CaM kinase II [13], PKA [13], PKC family kinases [14]) and XXSL, suggesting that the activities of members of PKC family kinases and PKA kinases were likely decreased in early-stage HCC (Figure 1F, Table S9).

We further performed kinase-substrate enrichment analysis (KSEA) to interpret the regulation of kinase activities in HCC. We focused on the kinases with significant phosphorylation regulation in early-stage HCC, and the KSEA analysis revealed seven kinases with significantly elevated activities in HCC (Figure 1G, Table S10), including several cyclin-dependent kinases (CDK1, CDK2, CDK7, CDK9, and CDK12) and cell cycle regulator WEE1, mitogenactivated protein kinase 6 (MAPK6), and dual specificity protein kinase CLK1. Four kinases showed decreased activity in early-stage HCC, including protein kinase C alpha (PRKCA), cAMP-dependent protein kinase catalytic subunit alpha (PRKACA), calcium/calmodulindependent protein kinase type II subunit delta (CAMK2D) and ribosomal protein S6 kinase alpha-1 (RPS6KA1) (Figure 1G). As indicated by the results of motif and KSEA analysis, further verification of the phosphorylation levels on activation sites of these kinases confirmed the activity elevation of CDK1, CDK2, CDK7, CDK9, CK1D, CK1E, SRPK1, and CLK1-4, as well as the decrease in the activity of PRKACA, PRKCA, and CAMK2D in Jiang et al.’s cohort [4] and Gao et al.’s cohort [7] (Figure S2). In summary, by integrating motif enrichment and KSEA analyses, we confirmed the elevated or decreased activities of these kinases, which likely play a major role in driving dysregulated protein phosphorylation in HCC.

### The EGFR-mediated canonical ERK/MAPK and atypical MAPK signaling cascades in early-stage HCC

The enrichment map of pathways that were dysregulated only in protein phosphorylation showed that several pathways, including the EGF pathway and MAPK family cascades, were observed with solely a decrease in phosphorylation levels (Figure 1E). Surprisingly, even though the majority of EGFR-mediated canonical ERK/MAPK signaling molecules, especially the MAPK1 and MAPK3, showed significantly elevated protein expression in early-stage HCC, phosphorylation regulation on the activation sites of the proteins in EGFR-mediated canonical ERK/MAPK signaling were significantly decreased in HCC (**Figure 2**A). In contrast, the phosphorylation regulation on the activation sites of RPS6KA3, together with MAPK6 (an atypical MAPK), was significantly elevated in HCC in both Jiang et al.’s cohort and Gao et al.’s cohort (Figure 2A, Figure S3A and B).

**Figure 2:**
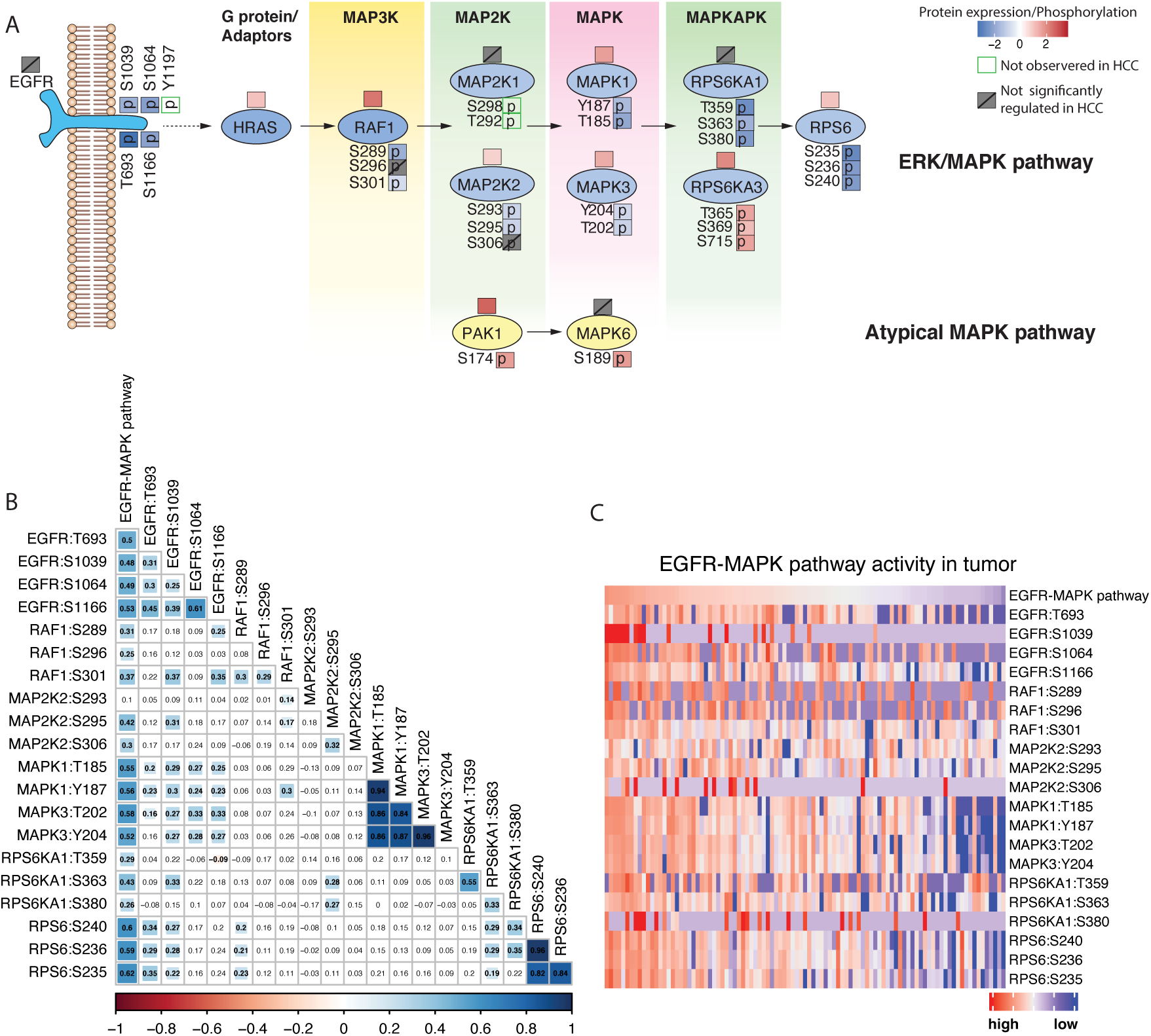
Expression and phosphorylation level of the EGFR-mediated canonical ERK/MAPK and atypical MAPK signaling cascades in HCC in Jiang et al.’s cohort. (A) The diagram of expression and functional phosphosite of proteins in the EGFR-mediated ERK/MAPK pathway and atypical MAPK pathway in HCC. The upper box on the protein diagram represents the expression level of the protein, while the lower box indicates the phosphorylation status of the specific site, marked with a ‘p’ inside the box. Phosphosites not detected were indicated by a blank green box, while proteins and phosphosites with no significant regulation in HCC were marked with a grey box containing a forward slash. (B) The correlation of the phosphorylation level of activation sites of proteins in EGFRmediated canonical ERK/MAPK signaling cascades in HCC. The phosphorylation levels of activation sites in proteins within the EGFR-mediated canonical ERK/MAPK signaling cascade in HCC were analyzed for correlations. Correlation coefficients are shown in squares, with significant ones highlighted in color. The median value of their activation sites was taken as a proxy of EGFR-mediated canonical ERK/MAPK pathway, and (C) patients were ranked from high to low by that, and the heatmap of their phosphorylation levels was shown (see also Figure S3).

Phosphorylation of the sites Thr693, Ser1039, Ser1064, Ser1166, and Tyr1197 on EGFR underwent increased regulation upon EGF stimulation [15]. The canonical ERK/MAPK signaling acts as a downstream effector of several tyrosine kinases including EGFR, and RPS6KA1 (p90RSK) lies at the terminus of the canonical ERK/MAPK pathway [16, 17]. Once activated, RPS6KA1 dissociates from MAPK1/MAPK3, translocates to the nucleus [16, 18], and phosphorylates RPS6 [19]. Phosphorylation of Ser235, Ser236, and Ser240 on RPS6 also increases upon EGF stimulation [15, 19]. Therefore, using the median of z-score-scaled phosphorylation levels of these phosphosites as a proxy for EGFR-mediated canonical ERK/MAPK pathway activity, which showed a significant positive correlation with the phosphorylation levels of the activation sites on the majority of molecules in this signaling (Figure 2B, Figure S3C). Furthermore, ranking patients from high to low based on EGFR-mediated canonical ERK/MAPK pathway activity revealed consistent regulation of this signaling cascade in HCC (Figure 2C, Figure S3D).

### Phosphoproteomic subtypes resolve the heterogeneity of early-stage HCC

PCA analysis showed the complex heterogeneity of the phosphoproteome in HCC (Figure 1A), and the dysregulation of signaling pathways mediated by phosphorylation also revealed distinct regulation of the phosphoproteome (Figure 1E, Figure 2), which prompted us to explore the molecular subtypes in the phosphoproteome of HCC. NMF factorization and consensus clustering of the phosphoproteome presented three subtypes from Jiang et al.’s cohort: the pS1 subtype (n=35), pS2 subtype (n=28), and pS3 subtype (n=34) (Figure S5 A-C, **Figure 3**A). Alignment of the proteomic subtypes [4] and clinical characteristics showed no significant difference in serum AFP status ( *≥* 200 ng ml*^−^*^1^), microvascular invasion (MVI^+^), and patient age among the three phosphoproteomic subtypes (Figure 3C and D). Interestingly, all female patients were distributed in the pS1 and pS2 subtypes, whereas the pS3 subtype included only male patients ( Fisher’s exact test, *p* = 0.0022). The pS1 subtype comprised the majority of patients in the proteomic SI subtype, low degree of MVI^+^ status, and a small tumor size (Figure 3C and D). The pS2 subtype was characterized by a larger tumor size than the pS1 subtype (Figure 3D). Patients in the pS3 subtype had a trend of a higher degree of MVI^+^ status and the worst prognosis in overall survival (Figure 3B and C). Notably, the pS3 subtype encompasses patients classified within the proteomic SI and SII subgroups, suggesting that early phosphorylation alterations have been detectable in these samples.

**Figure 3:**
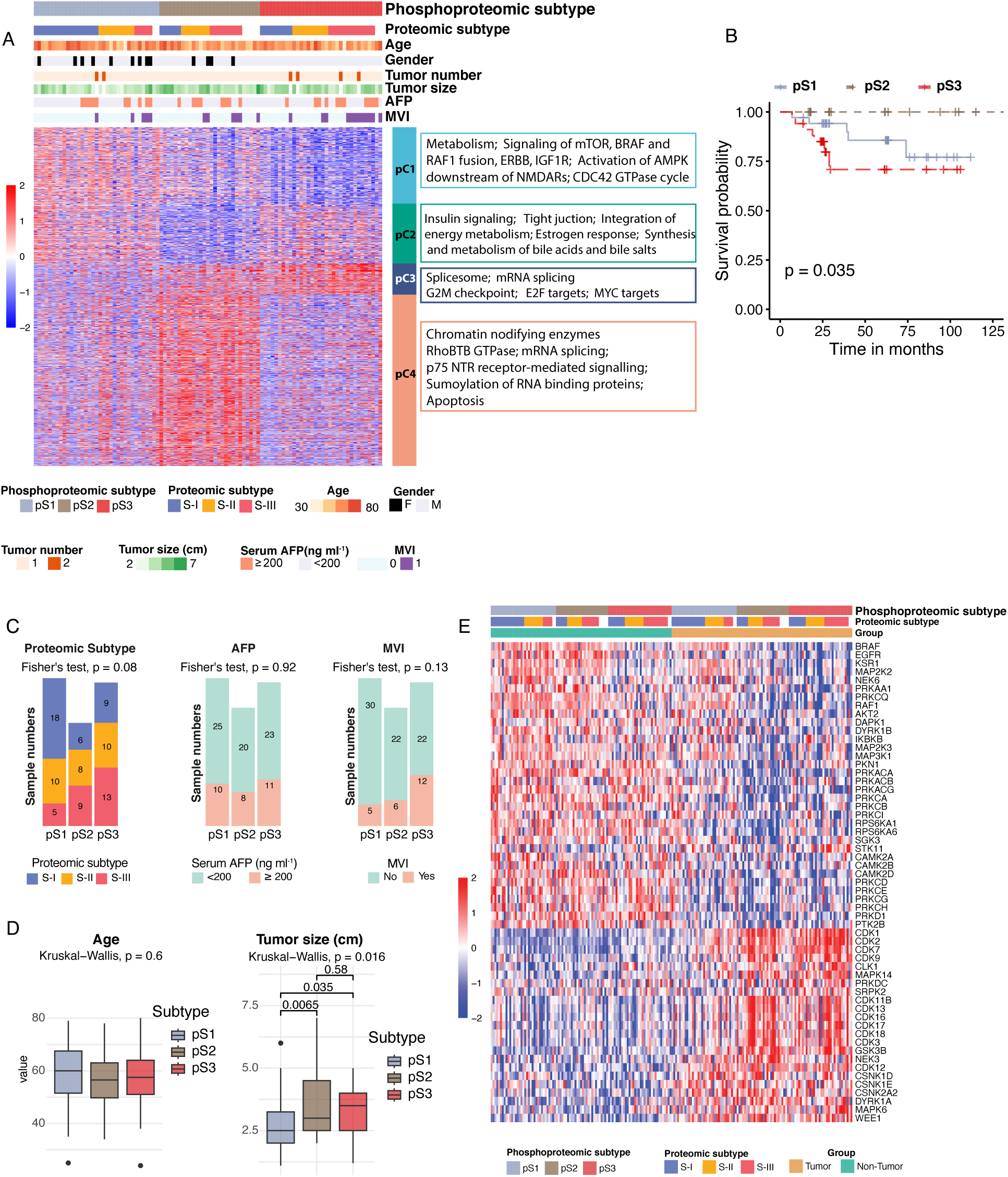
(previous page.) **Phosphoproteomic subtypes identified in HCC tissues and their correlation with clinical characteristics and kinase activities** (A) Nonnegative matrix factorization and consensus-clustering analysis of the phosphoproteome in HCC tissues stratified patients into three phosphoproteomic subtypes. The proteomic subtypes and clinical characteristics from Jiang et al.’s cohort were annotated in the column annotation. A total of 1,816 phosphosites showed significantly differential levels among the three subtypes and were divided into four clusters. The relative abundance of phosphosites (log2-transformed) was shown in the heatmap. Specifically, the pC1 cluster: phosphosites with the highest abundance in the pS1 subtype. pC2 cluster: phosphosites exhibiting the lowest abundance in the pS2 subtype. pC3 cluster: phosphosites showing the highest abundance in the pS3 subtype. pC4 cluster: phosphosites with the highest abundance in the pS2 subtype. (B) Kaplan–Meier curves for overall survival (OS), based on the phosphoproteomic subtypes (log-rank test). (C) Association of phosphoproteomic subtypes with proteomic subtypes and clinical features, including serum AFP levels, MVI, (D) patient age, and tumor size. (E) Kinases inferred with significantly different activities in the phosphoproteomic subtypes in HCC, shown together with the corresponding non-tumor samples. Statistical information: the box plots (D) show the median (center line), interquartile range (box bounds, 25th–75th percentiles), and whiskers extending to the largest and smallest values within 1.5 *× IQR* from the hinges. Data beyond the whiskers are plotted as individual points. The exact sample sizes of pS1, pS2, and pS3 are *n*=35, 28, 34 for panels B and D, respectively.

We identified 1,816 phosphosites that were significantly regulated among the phosphoproteomic subtypes and categorized them into four clusters (Figure 3A, Table S11): pC1-pC4, with which a clear separation of the three phosphoproteomic subtypes was presented (Figure S5D). To validate the robustness of the phosphoproteomic subtypes, we applied an SVM classifier (Figure S6A) trained on a balanced panel of 512 of the 1,816 phosphisites to the Gao et al.’s cohort, which successfully stratified the patients (log-rank *p* = 0.002, FigureS6) and the pS3 subtype consistently exhibited the aggressive clinical characteristics, including the poorest overall survival, significantly larger tumor sizes, and a higher proportion of elevated preoperative AFP compared to the pS1 subtype (Figure S6C and E). Assignment of the phosphoproteomic subtypes onto the proteome identified 566 proteins that were differentially expressed among the phosphoproteomic subtypes, which were also divided into four clusters: C1-C4 (Figure S8A). The pS1 subtype is characterized with higher abundance of phosphosites on proteins involved in metabolism of carbohydrates (ChREBP pathway), fatty acid, organic acid, vitamins, and the catabolism of branched chain amino acids (Figure S7B and C, Table S12) in tumor tissues. Meanwhile, the higher abundance of metabolism-related pathways was also observed on proteome in pS1 (Figure S8B and C, Table S14). In addition, signaling transduction with higher phosphorylation regulation in pS1 subtype included mTOR, BRAF and RAF1 fusion, ERBB, IGF1R, activation of AMPK downstream of NMDARs, CDC42 GTPase cycle and CREB signaling, etc. ( Figure S7D, Table S12), among these included the cascades on EGFR-mediated canonical ERK/MAPK signaling, i.e., Ser1064 and Ser1166 on EGFR, Thr202 and Tyr204 on MAPK3, and Thr185 and Tyr187 on MAPK1 (Figure S7A). The pS2 subtype presented significantly low abundance of phosphosites on proteins involved in the insulin signaling pathway, tight junction, PPARA and keratinocyte pathway, estrogen response, integration of energy metabolism, EIF4 pathway, synthesis and metabolism of bile acids and bile salts etc, a correlation of dysregulation of which and HBV-related HCC had been recently reported [7] (Figure 3A , Figure S7B and D). Besides, lower protein expression in the metabolism of carbohydrates, amino acids and derivatives, fatty acids etc. was observed in the pS2 subtype (Figure S8B and C). On other side, the pS2 subtype showed a distinctly high abundance of phosphosites on proteins (pC4 cluster) participating in signaling pathways related to RhoBTB GTPase, mRNA splicing, p75 NTR receptor-mediated signaling, apoptosis, chromatin-modifying enzymes, sumoylation of RNA binding proteins (Figure 3A, Figure S7B and D), suggesting regulation on transcription and epigenetic changes may have occurred in the pS2 subtype, which was also confirmed in the biological processes (Figure S7C, Table S13). As expected, the proteins with higher abundance in the pS2 subtype were enriched in epigenetic regulation of gene expression, basal transcription factors, rRNA processing and modification, transcription regulation by polymerase I, II, III, and hallmark of MYC targets, etc. (Figure S8, Table S14, S15). The pS3 subtype is characterized by a lower abundance of phosphosites on proteins that participate in metabolism, especially liver function-related pathways. Additionally, pS3 subtype showed high abundance of phosphosites on proteins involved in mRNA splicing, G2M checkpoint, and hallmarks of E2F and MYC targets (Figure 3A ), mRNA processing and spliceosome, base excision repair, and cellular component disassembly (Figure S7B and D).

Moreover, the pS2 and pS3 subtypes presented regulation on distinct phosphosites of proteins in the same pathways. For instance, the phosphorylation of Ser40 on MCM2, a marker of DNA replication during the cell cycle [20], was hyperphosphorylated in the pS2 subtype (Figure S7A). Whereas Ser27 on MCM2, phosphorylated by CDKs, showed a higher phosphorylation level in both the pS2 and pS3 subtypes (Figure S7A), suggesting proliferative characteristics mediated by distinct signaling in the pS2 and pS3 subtypes. Interestingly, proteins with high abundance in the pS3 subtype were involved in extracellular matrix organization (ECM), collagen formation, hallmarks of EMT, and angiogenesis (Figure S8B and C). EMT progression is associated with angiogenesis, and HCC cells acquire a more invasive and aggressive phenotype when they undergo this characteristic [21], implying a role of ECM and EMT in the progression of the pS3 subtype.

Blocking aberrant kinase activities is one of the strategies for anticancer therapeutics; therefore, we inferred kinase activities by single-sample kinase-substrate relation set enrichment analysis (ssKSEA). We found that the activities of 57 kinases were significantly regulated in phosphoproteomic subtypes (Figure 3E, Table S16). Notably, EGFR and canonical ERK/MAPK signaling, including MAP2K2 and RAF1, showed low activity in tumors, especially in the pS3 subtype, which is consistent with our previous analysis. In addition, members of the protein kinase A and the protein kinase C families showed declined activities in pS3, with even greater declines in pS2. Meanwhile, CDKs, MAPK14, and GSK3B showed not only elevated activities in tumors, but also even greater increases in pS2 and pS3 subtypes. We noticed that CLK1 and SRPK2, two kinases that contribute to the phosphorylation of splicing factors, presented higher activities in the pS3 subtype. Taken together, the integrated analysis of the phosphoproteome and proteome uncovered distinct phosphoproteomic subtypes in early-stage HCC, characterized by unique molecular signatures, clinical features, signaling pathways, and kinase activities among HCC patients.

### SRSF3 contributes to the proliferation, migration, and invasion of HCC cells

We then focused on the pS3 subtype, which was characterized by the worst prognosis, EMT phenomenon, and hyperphosphorylation regulation of proteins involved in the spliceosome and mRNA processing (Figure 3B). Among the phosphosites in pC3 that showed high levels in the pS3 subtype, these were involved in phosphosites on spliceosome components such as SRRM2, PRPF3, and splicing factor SRSF3. Overexpression of SRSF3 has been shown to contribute to the tumor progression of glioblastoma (GBM), head and neck squamous cell carcinoma (HNSCC) [22], and promotes gemcitabine resistance in pancreatic cancer [23]. SRSF3 is crucial for the functional differentiation of hepatocytes [24]; however, decreased levels of SRSF3 have been reported in HCC, and its loss in hepatocytes promoted the development of metastatic HCC in mice [25]. We found SRSF3 was significantly upregulated in tumor tissues in Jiang et al.’s cohort and Gao et al.’s cohort (**Figure 4**A), and this was further confirmed by immunohistochemistry (IHC) staining analysis (Figure 4B and C). Therefore, we hypothesized that the elevation of SRSF3 may play a crucial role in the carcinogenesis and development of HCC. Stable SRSF3 knockdown using short hairpin RNA (shRNA) in MHCC-97H and Huh-7 cell lines significantly inhibited cell proliferation (Figure 4D and E), cell monoclonal formation (Figure 4F ), and reduced the ability of migration and invasion of HCC cells (Figure 4G). Conversely, stable overexpression of SRSF3 in PLC/PRF/5 and MHCC-97H cell lines significantly promoted the malignancy of HCC cells (Figure S9A to D) in both cell lines. These results indicated that SRSF3 contributes to the proliferation, migration, and invasion of HCC cells.

**Figure 4:**
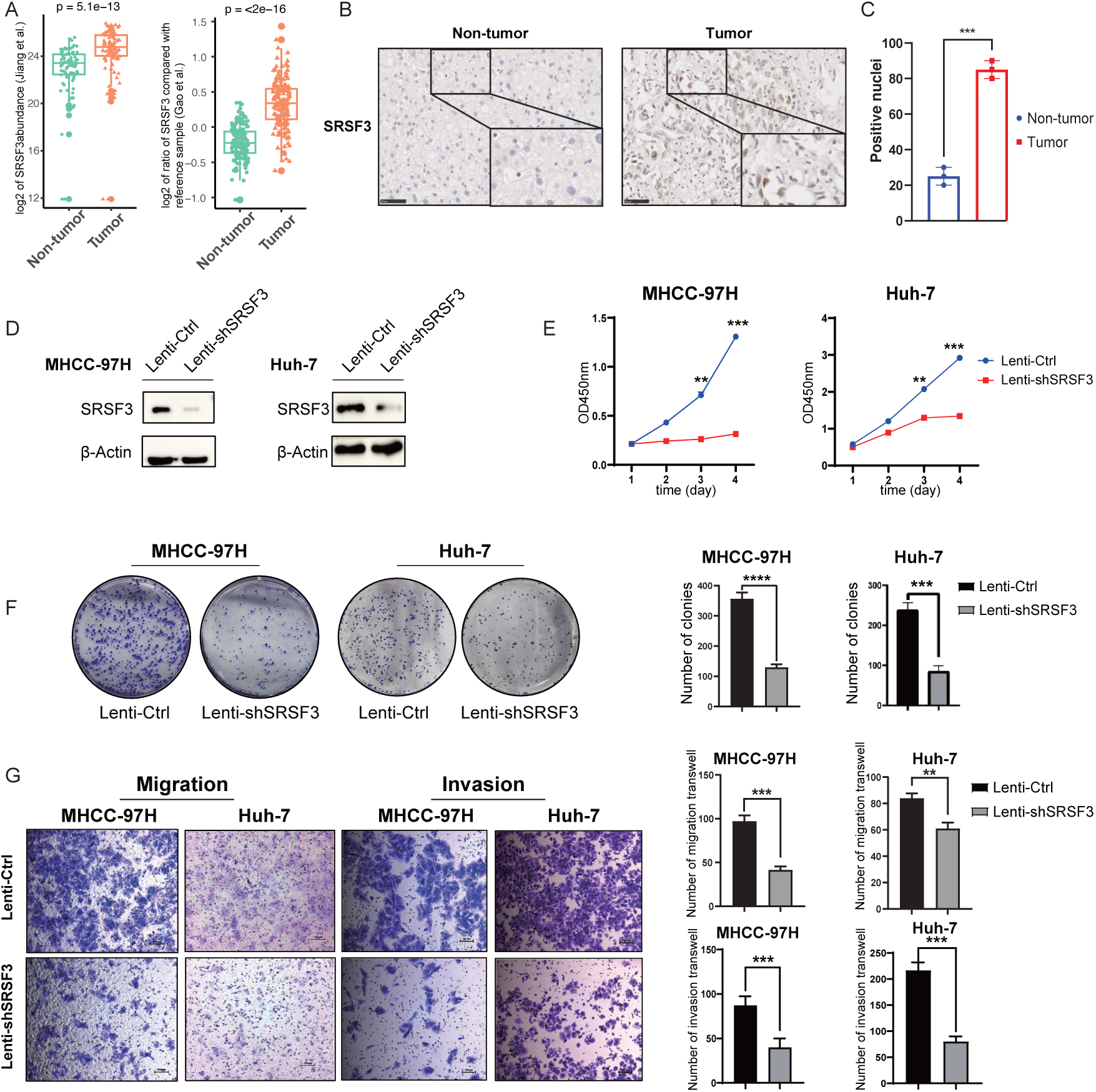
SRSF3 was upregulated in HCC, and stable SRSF3 knockdown in HCC cell lines significantly decreased the malignancy of HCC cells. (A) SRSF3 was significantly upregulated in tumor tissues of HCC patients in Jiang et al.’s cohort and Gao et al.’s cohort. (B) Digital scans of representative IHC staining of SRSF3 and (C) statistical analysis of the positive nuclei in the non-tumor and tumor tissues of HCC patients. (D) Stable SRSF3 knockdown in MHCC-97H and Huh-7 HCC cell lines significantly reduced the capabilities of (E) cell proliferation, (F) cell clone formation, (G)migration, and invasion. Statistical information: the box plots (A) definitions are as in Figure 3. Bar graphs (C, F, G) are mean *±* SD; the exact sample size is *n*=3 for panels C, F, and G. For all graphs, asterisks show statistical significance by two-sided Wilcoxon rank-sum test, \**p <*0.05; \*\**p <*0.01; \*\*\**p <*0.001.

### SRSF3 hyperphosphorylation promotes migration, invasion and proliferation of HCC cells

Integrative phosphoproteomic analysis revealed that phosphorylation of SRSF3, rather than SRSF3 protein expression, was significantly upregulated in the pS3 subtype (**Figure 5**A). These residues are localized in the domain rich in arginine and serine (RS domain) at the C-terminus of SRSF3, and reversible phosphorylation on the RS domain of the SRSF family was found to regulate their activities in spliceosome maturation, splicing catalysis and mRNA export [26, 27]. To identify the functional phosphosites driving the pS3 phenotype, we examined the phosphorylation patterns of SRSF3. The phosphoproteomic data revealed that Ser148 and Ser150 showed striking quantitative synchronization in the samples (Figure S10A). Crucially, we observed that Ser148 and Ser150 showed an extremely high quantification correlation across the tumor samples (r= 0.95, Figure S10B) and identical quantification intensity values in 83.5% of tumor cases (*n*=97, Figure S10A).

**Figure 5:**
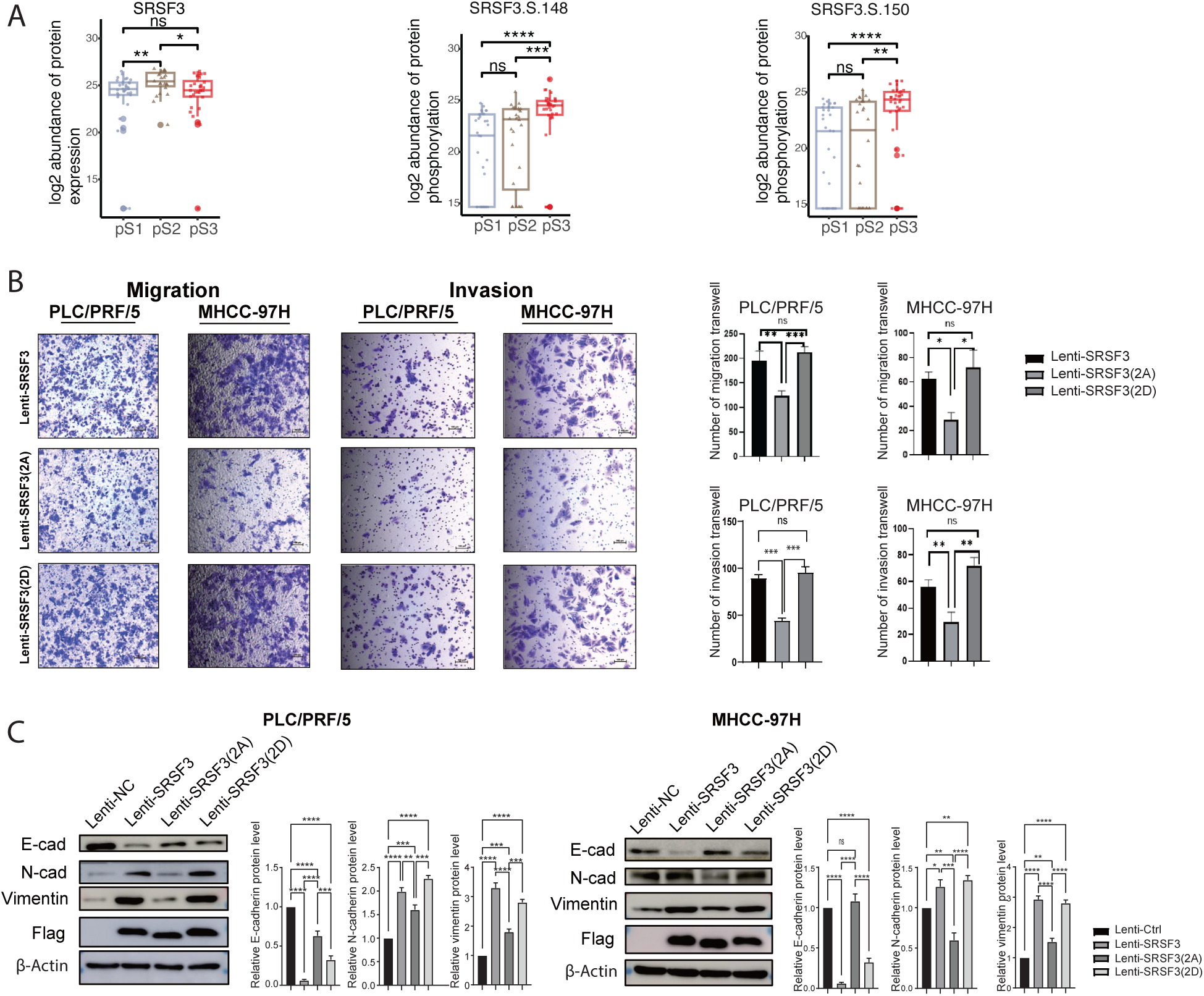
Phosphorylation on Ser148 and Ser150 of SRSF3 promotes migration and invasion in HCC cells. (A) The phosphorylation levels on Ser148 and Ser150 of SRSF3 were elevated in the phosphoproteomic pS3 subtype, while no elevation of the protein expression of SRSF3 was observed. (B) Overexpression of the phosphorylated mutant (S148D/S150D) and the wild-type SRSF3 (SRSF3-WT) significantly increased migration and invasion capabilities compared to the dephosphorylated mutant (S148A/S150A) in HCC cells. (C) Overexpression of phosphorylated mutant (S148D/S150D) and the SRSF3-WT promoted epithelial-mesenchymal transition (EMT) of HCC cells, while the dephosphorylated mutant (S148A/S150A) showed a limited effect on the promotion of EMT compared to the phosphorylated mutant. Statistical information: the box plot and error bar definitions are as in Figures 3 and 4. The sample sizes in panel A are as in Figure 3D, *n*=3 for panels B and C. For all graphs, asterisks show statistical significance based on unadjusted *p*-values by two-sided Wilcoxon rank-sum test, \**p <*0.05; \*\**p <*0.01; \*\*\**p <*0.001.

The pS3 subtype is also characterized by EMT; therefore, we hypothesized that elevated phosphorylation levels at Ser148 and Ser150 on SRSF3 in the pS3 subtype of HCC (Figure 5A) may be an important factor affecting the prognosis of HCC patients. The striking co-regulation of Ser148 and Ser150 suggested them as a regulatory unit distinct from the more variable adjacent site Ser152. Therefore, we focused on the Ser148/Ser150 doublet for functional mutagenesis. We constructed stable cell lines expressing SRSF3 with mutations simulating the dephosphorylated (S148A, S150A), phosphorylated (S148D, S150D), and wild-type forms (Figure S11A). The construction of the cell lines was verified by western blot analysis (Figure S11B). As expected, overexpression of the phosphorylated mutant (S148D/S150D) and SRSF3-WT in PLC/PRF/5 and MHCC-97H cell lines showed significantly increased migration and invasion capabilities compared to the dephosphorylated mutant (S148A/S150A) in HCC cells (Figure 5B), accompanying by increased expression of mesenchymal markers N-cadherin and vimentin, and decreased expression of epithelial marker E-cadherin, suggesting the promotion of EMT in HCC cells; whereas the dephosphorylated mutant (S148A/S150A) had little effect on promotion of EMT (Figure 5C). In addition, the phosphorylated mutant (S148D/S150D) and the SRSF3-WT form both showed significantly increased abilities for cell monoclonal formation and cell proliferation compared to the dephosphorylated mutant (S148A/S150A) (Figure S11C and D). Overexpression of SRSF3-WT resulted in a significant increase in tumor weight compared with the control group, whereas the SRSF3-S148D/S150D mutant resulted in a significant increase in tumor weight compared to the SRSF3-S148A/S150A mutant in the PLC/PRF/5 cell line xenograft model (Figure S11E and F). Interestingly, the SRSF3 S148D/S150D mutant showed a significant increase in xenograft growth compared to the SRSF3-WT form (Figure S11G). Ki-67, a prominent proliferation marker used in pathology [28], not only showed significantly elevated expression in the SRSF3-S148D/S150D mutant xenograft compared to the SRSF3-S148A/S150A mutant xenograft, but also showed higher expression compared to the SRSF-WT xenograft (Figure S11H and I), suggesting a higher proliferative ability of HCC cells mediated by phosphorylation regulation of Ser148 and Ser150 on SRSF3. Taken together, these results indicate that higher phosphorylation levels of SRSF3 at Ser148 and Ser150 promote EMT, metastatic invasion, and proliferation in HCC cells.

### Activated SRPK1 and CLKs mediate phosphorylation of Ser148 and Ser150 at SRSF3 and inhibitors targeting SRPK1 or CLKs suppress hyperphosphorylation of SRSF3-mediated migration and invasion in HCC

SRPK1, SRPK2, and members of the CLK family act as upstream kinases of the SRSF family [29, 30] and cooperate tightly to regulate the function of SRSF proteins [31]. Phosphorylation of SRPK1 at Ser309 and Ser311 was significantly elevated in HCC (Figure S2A and B), with Ser309 serving as a self-activation site for the kinase [32], indicating that SRPK1 was activated in HCC. Although the SRPK2 kinase protein was significantly upregulated in HCC (Table S5), the phosphorylation levels of its kinase activation sites, Ser494 and Ser497 [33], the former was significantly downregulated in HCC, whereas the latter showed inconsistent trends between Jiang et al’s cohort and Gao et al’s cohort (Figure S2A and B). Even though CLK1-4 kinases (CLKs) were not detected at the protein level, phosphorylation levels of activation sites on CLKs [34] were all significantly increased in HCC (Figure S2). Given the elevated kinase activities of SRPK1 and CLKs in HCC (Figure S2A and B), we propose that they may be the upstream kinases that phosphorylate SRSF3 in HCC. We then treated the PLC/PRF/5 and MHCC-97H cells with SRPK1 inhibitor SRPIN340 [35] and CLKs inhibitor TG003 [36] in each cell line, respectively. Both SRPIN340 and TG003 effectively reduced the phosphorylation levels of Ser148 and Ser150 on SRSF3 in PLC/PRF/5 and MHCC-97H cell models (**Figure 6**A, B, and S12E), confirming that the phosphorylation of Ser148 and Ser150 on SRSF3 is regulated by SRPK1 and CLKs in HCC. We then treated PLC/PRF/5 and MHCC-97H cells that stably overexpress SRSF3-WT with individual inhibitors, and the migration and invasion ability of SRSF3-WT cells significantly decreased to a level similar to that of the dephosphorylated mutant SRSF3 (S148A/S150A) (Figure 6C). Moreover, under treatment with SRPIN340 or TG003, the SRSF3-WT showed significantly decreased abilities in migration and invasion in comparison with the phosphorylated mutant SRSF3 (S148D/S150D) (Figure S12A). Taken together, these findings indicate that SRPK1 and CLKs phosphorylate SRSF3 at Ser148 and Ser150, thereby promoting the migration and invasion of HCC cells. Consequently, targeting these kinases may offer a promising strategy for the development of targeted therapies for HCC patients.

**Figure 6:**
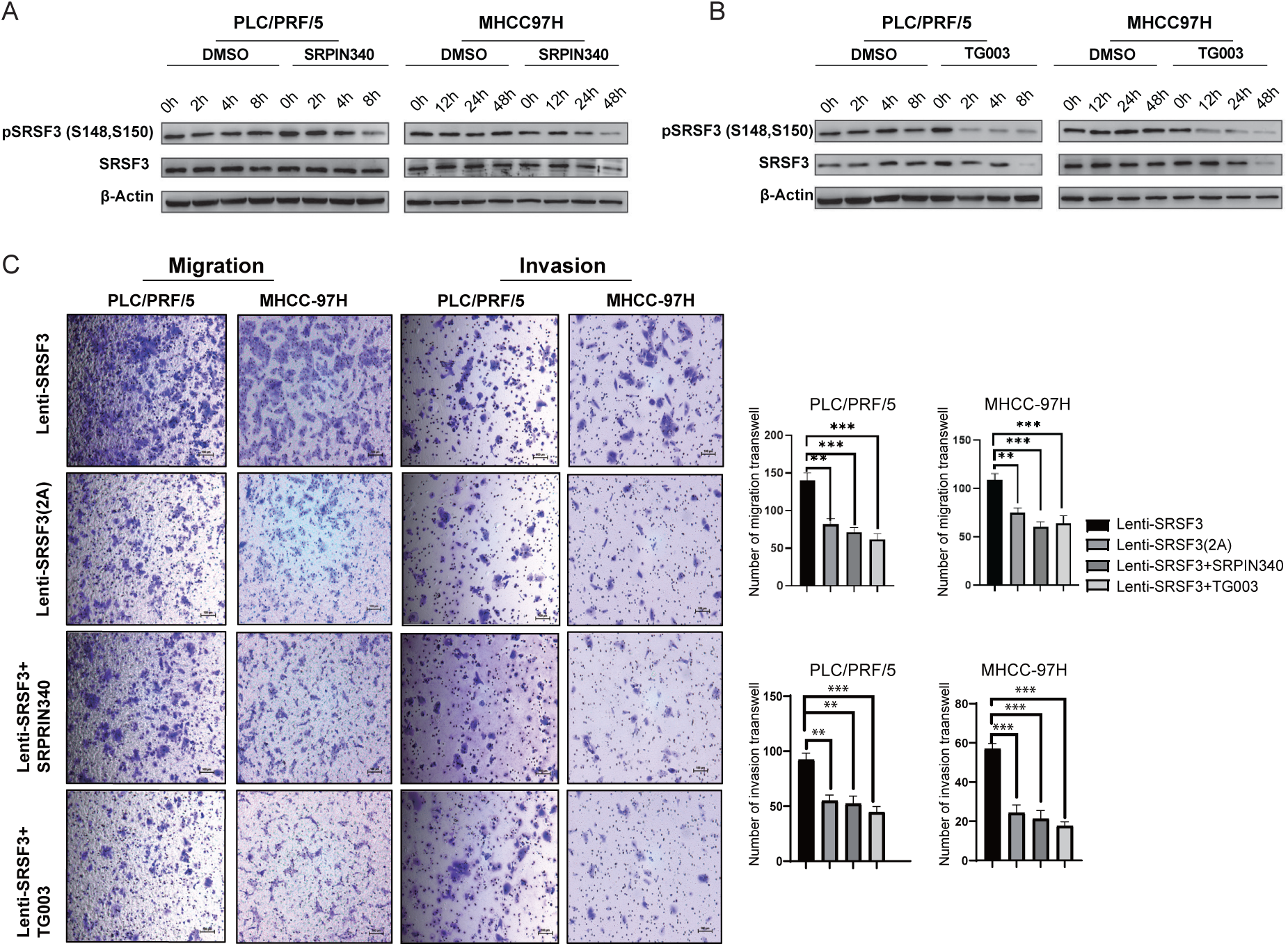
SRPK1/CLKs phosphorylate Ser148 and Ser150 on SRSF3, and inhibitors targeting SRPK1 or CLKs significantly suppress migration and invasion of HCC cells. (A) SRPIN340 and (B) TG003 were applied to PLC/PRF/5 cells (50 *µ*M) and MHCC-97H cells (100 *µ*M) for the indicated times (see also Figure S12D). Addition of (A) the SRPK1 kinase inhibitor SRPIN340 or (B) the CLKs kinase inhibitor TG003 effectively reduced the abundance of phosphorylation at Ser148 and Ser150 on SRSF3 in these cell models in a time-dependent manner. (C) Treatment with SRPIN340 or TG003 to SRSF3-WT overexpressing PLC/PRF/5 and MHCC-97H cells significantly reduced the migration and invasion capabilities of HCC cells. The dephosphorylated mutant SRSF3(S148A/S150A), without treatment with SRPIN340 or TG003, also showed decreased migration and invasion capabilities compared to the SRSF-WT. Statistical information: the error bar definition is as in Figure 4, *n*=3 for panels C. For all graphs, asterisks show statistical significance by two-sided Wilcoxon rank-sum test, \**p <*0.05; \*\**p <*0.01; \*\*\**p <*0.001.

### Hyperphosphorylation of SRSF3 in the pS3 subtype is potentially mediated through suppression of PP1 phosphatase activity via phosphorylation on PPP1R7

Given that phosphorylation on Ser148 and Ser150 on SRSF3 is significantly elevated in the pS3 subtype (Figure 5A), as well as that SRPK1 and CLKs contribute to phosphorylation of Ser148 and Ser150 on SRSF3, we speculate that phosphorylation of the activation sites of these kinases would have been significantly elevated in the pS3 subtype of early-stage HCC. Surprisingly, neither the protein expression nor the phosphorylation of active sites of these kinases was observed with elevation in pS3 subtype (Figure S12B and C), suggesting the regulation of SRSF3 by phosphatases may be inhibited in the pS3 subtype. Protein phosphatase 1 (PP1) mediates dephosphorylation in the RS domain of SRSF1 [37], PPP1R7 (Sds22) is a conserved PP1 regulator and a PP1 trap [38], and phosphorylation of the N-terminal of PPP1R7 inhibits PP1 activity [39]. We found that phosphorylation levels at Ser12, Ser24, and Ser27 of PPP1R7 showed high abundance in the pS3 subtype (Figure S12C); suggesting that PP1 phosphatase activity may be suppressed in this subtype, thereby leading to increased SRSF3 phosphorylation. As expected, the treatment of HCC cells with Tautomycetin, a PP1 phosphatase inhibitor, significantly increased the phosphorylation levels of Ser148 and Ser150 on SRSF3 (Figure S12G), confirming that PP1 regulates the dephosphorylation of SRSF3 in HCC. Taken together, we have demonstrated that activation of SRPK1 and CLK1-4, potentially coordinated with suppression of PP1 phosphatase activity by phosphorylation of PPP1R7, determined elevation of phosphorylation of SRSF3 in the pS3 subtype, thereby contributing to the worse prognosis of HCC patients in the pS3 subtype.

## Discussion

HCC is one of the most challenging cancers, characterized by high recurrence and mortality, and the nearly identical incidence and mortality underscore the dismal prognosis of this disease [1, 40, 41], which calls for an urgent need for a systematic understanding of its molecular landscape. PTMs dramatically increase the diversity of protein functions; however, previous studies have seldom investigated the specific regulation of protein phosphorylation on a large scale in HCC, and the information on the dynamic regulation of phosphosites was lost when considering them with collapsed phosphoproteins [5, 7]. In this study, protein phosphorylation and expression levels showed a weak positive correlation in early-stage HCC, suggesting the role of post-translational regulation by phosphorylation in cellular signaling (Figure 1B and C). Integration of the phosphoproteome with the proteome in early-stage HCC revealed unique regulation observed only at the level of protein phosphorylation, such as transcription regulated by FOXO, NOTCH1, and RORA (Figure 1E), which may be involved in carcinogenesis and development of HCC.

Persistent activation of Ras and downstream ERK in canonical ERK/MAPK signaling cascades has been previously reported in HCC [42]. Conversely, in this study, the phosphorylation levels (rather than protein expression) of EGFR-mediated canonical ERK/MAPK signaling cascades, together with PRKCA, PRKACA, and CAMK2D, were significantly downregulated in HCC. The downregulation of phosphorylation of EGFR-ERK/MAPK likely reflects profound negative feedback loops that are hyper-activated in specific HCC contexts. For instance, hypoxia-inducible factors in tumors can upregulate dual-specificity phosphatases (DUSPs) that dephosphorylate and inactivate ERK/MAPK signaling [43]. Furthermore, microenvironmental stress frequently triggers aberrant endocytosis and intracellular trafficking of EGFR [44], which is thought to initiate termination of the MAPK signaling cascade [45]. To further confirm the uncoupling of phosphorylation from protein expression in EGFRERK/MAPK, we performed the site-level residual analysis (regressing out total protein) and generated patient-ranked heatmaps (Figure S4, Table S17). These analysis demonstrated that the active suppression and heterogeneity of EGFR-ERK/MAPK phosphorylation are not fully determined by protein abundance in the tumor cohorts. The canonical ERK/MAPK signaling cascades act as downstream of several tyrosine kinases, including the drug targets of sorafenib and lenvatinib [46]. The significant downregulation of EGFR-mediated MAPK signaling in HCC may explain the low objective response rate (ORR) of sorafenib and lenvatinib in HCC patients. Moreover, the activities of atypical MAPK signaling molecules such as MAPK6, RPS6KA3, CDKs, CLKs, and SRPK1 were significantly upregulated in HCC (Figure 2B and S3A), suggesting the potential therapeutic benefits of targeting these kinases in HCC.

Interrogating the heterogeneity of the phosphoproteome uncovered three phosphoproteomic subtypes of HCC, each characterized by distinct phosphosite signatures, signaling pathways, and kinase activities. The pS3 subtype has the poorest prognosis . Furthermore, the exclusive male composition in the pS3 subtype perfectly aligns with the established molecular roles of sex steroid receptors in HCC [47]. The androgen receptor signaling has been suggested to promote HCC [48], whereas estrogen receptor signaling elicits a protective effect [49]. Therefore, the pS3 subtype likely represents an aggressive clinical state related to these underlying differences in sex steroid receptor signaling. Activation of the EMT program is capable of promoting the dissemination of tumor cells during the early stages of tumorigenesis [2]. The pS3 subtype exhibits a strong EMT signature on the proteome (Figure S8C), suggesting a potential for early-stage dissemination. Furthermore, our *in vitro* functional assays suggest that EMT-associated processes may be enhanced through SRSF3 hyperphosphorylation at Ser148 and Ser150, which is regulated by SRPK1 and CLKs in HCC. Several CDKs, WEE1, MAPK6, CK1, and CK2, etc., exhibit significantly higher activity across all three phosphoproteomic subtypes compared to non-tumor tissues. Therefore, targeting the kinases SRPK1/CLKs and kinases active across all three phosphoproteomic subtypes could expand therapeutic options for early-stage HCC, offering a promising strategy to prevent progression to advanced disease.

SRSF3 shuttles between the nucleus and cytoplasm and requires phosphorylation regulation of the RS domain [50]. We then performed immunofluorescence experiments to examine whether different phosphorylation mutants of SRSF3 influenced its subcellular localization. The results showed that all of SRSF3-WT, SRSF3-S148A/S150A, and SRSF3-S148D/S150D localized in the nuclear speckles and nucleoplasm of PLC/PRF/5 cells (Figure S12G), suggesting that EMT may be mediated by other functions of SRSF3, such as mRNA export, mRNA stability, or regulation of translation [51], which deserves further investigation. Although the phosphorylation of Ser152 is also upregulated in the tumor group, and showed the highest expression in pS3 tumor subtype (Table S11), its phosphorylation showed a distinct quantification pattern compared with Ser148 and Ser150 (Figure S10A). This variance may be attributed to the unique structural biology of SRSF3. SRSF3 lacks a pseudo-RNA recognition motif (pseudo-RRM), making its RS domain easily exposed and more susceptible to dephosphorylation by phosphatases [52]. Moreover, while the triple phosphorylation Ser148/Ser150/Ser152 exists in a subset of patients, the Ser148/Ser150 phosphorylation covers a larger proportion of patients, and the most malignant pS3 subtype showed a higher proportion of Ser148 and Ser150 co-phosphorylation pattern (Figure S10C). Consequently, this stable Ser148/Ser150 phosphorylation acts as a crucial regulatory core required to promote the metastatic EMT phenotype.

In summary, we interrogated the phosphoproteome and proteome in early-stage HCC, enhancing our understanding of HCC heterogeneity at an early stage, uncovering the dysregulated phosphorylation signatures, kinases, and pathways in early-stage HCC and across phosphoproteomic subtypes, and demonstrated that SRSF3 hyperphosphorylation mediates the mechanism for development and metastasis of HCC, which should facilitate the development of proteomics-driven precision medicine for HCC patients.

## Materials and Methods

### Patient cohorts

The phosphoproteomic dataset with 27,651 phosphosites and the proteomic dataset with 9,254 proteins from 103 tumor and non-tumor paired samples were obtained from Jiang et al.’s cohort [4]. The phosphoproteomic dataset with 26,418 phosphosites and the proteomic dataset with 6,478 proteins from 159 paired tumor and non-tumor samples from Gao et al.’s cohort [7] were used to verify the findings of this study. Detailed methods are presented in the Supporting Methods section.

### Integrative phosphoproteomic and proteomic data analysis

We performed analysis on correlations of protein expression and phosphorylation level, dysregulated protein phosphorylation, and identification and characterization of phosphoproteomic subtypes in early-stage HCC. The detailed methods are presented in the Supporting Methods.

### Cell culture

The PLC/PRF/5, MHCC-97H, and Huh7 cell lines were cultured in DMEM/MEM medium containing 10% fetal bovine serum (FBS). The cells were maintained at 37°C and 5% CO2 in a cell culture incubator.

### Immunohistochemistry (IHC)

The required tissue sections were retrieved from the oven and immersed in xylene twice for 20 minutes each. They were then sequentially transferred through 100%, 100%, 95%, 95%, 90%, and 80% ethanol for 5 minutes each, followed by washing with 1 TBS for 5 minutes. Antigen retrieval was performed by citrate acid treatment, followed by blocking with 5% BSA and incubation with the primary antibody overnight. After adding the enhancement solution and secondary antibody, the sections were washed thrice with TBS, stained with DAB, counterstained with hematoxylin, dehydrated, mounted, and scanned. Sections were differentiated using hydrochloric alcohol and blued under running water.

### Statistical analysis

The data provided in this study were statistically analyzed in the R environment or using GraphPad Prism 9 software and presented as mean *±* standard deviation (mean *±* SD). Two-sample comparisons were performed using the two-sided Wilcoxon rank-sum test, with *p*-values *<*0.05 were considered significant. Associations between phosphoproteomic subtypes and clinical characteristics were evaluated using Fisher’s exact test for categorical data and the Wilcoxon rank-sum test for continuous data. Log-rank test and Kaplan–Meier survival curves were applied to compare the overall survival among the phosphoproteomic subtypes, and a significance level of 0.05 was used as the cut-off threshold. \**p <*0.05; \*\**p <*0.01; \*\*\**p <*0.001; \*\*\*\**p <*0.0001. The details and description of the statistical analysis methods are provided in the Supporting Methods.

## Ethical statement

Animal care and experiments were approved by the Institutional Animal Care and Use Committee (IACUC) of the National Center for Protein Sciences, Beijing, China (NCPSB) (Approval No. IACUC-20220524-32MT).

## Data Availability

No new primary data were generated in this study. The database accession numbers and repository links for the previously published datasets analyzed in this work are detailed in the Supporting Methods.

## Code Availability

All custom codes used for the analyses included in our manuscript were uploaded to the Bitbucket repository https://bitbucket.org/jiangbiolab/hccphosphostudy.

## CRediT author statement

**Liangliang Ren:** Conceptualization, Formal analysis, Investigation, Writing – original draft, Writing – review & editing. **Jiahan Jiang:** Investigation, Validation, Writing – original draft, Writing – review & editing. **Changwen Ning:** Validation, Writing – review & editing. **Weiran Lin:** Investigation, Writing – review & editing. **Lihui Qi:** Investigation, Writing – review & editing. **Chaoying Li:** Conceptualization, Supervision, Writing – original draft, Writing – review & editing. **Ying Jiang:** Conceptualization, Supervision, Writing – original draft, Writing – review & editing, Funding acquisition.

## Acknowledgements

This work was supported by the National Key R&D Program of China (Nos. 2024ZD0520500, 2021YFA1301601, 2020YFE0202200), the National Natural Science Foundation of China (Nos. 82273243, 32400596, 82090051, 92168207), and the Program of the State Key Laboratory of Proteomics (SKLP-K202405; SKLP-X202402; SKLP-O202401; SKLP-O202512).

## Supplementary Material

### Supplementary files

**File S1** Supporting Methods.

### Supplementary figures

**Figure S1.**
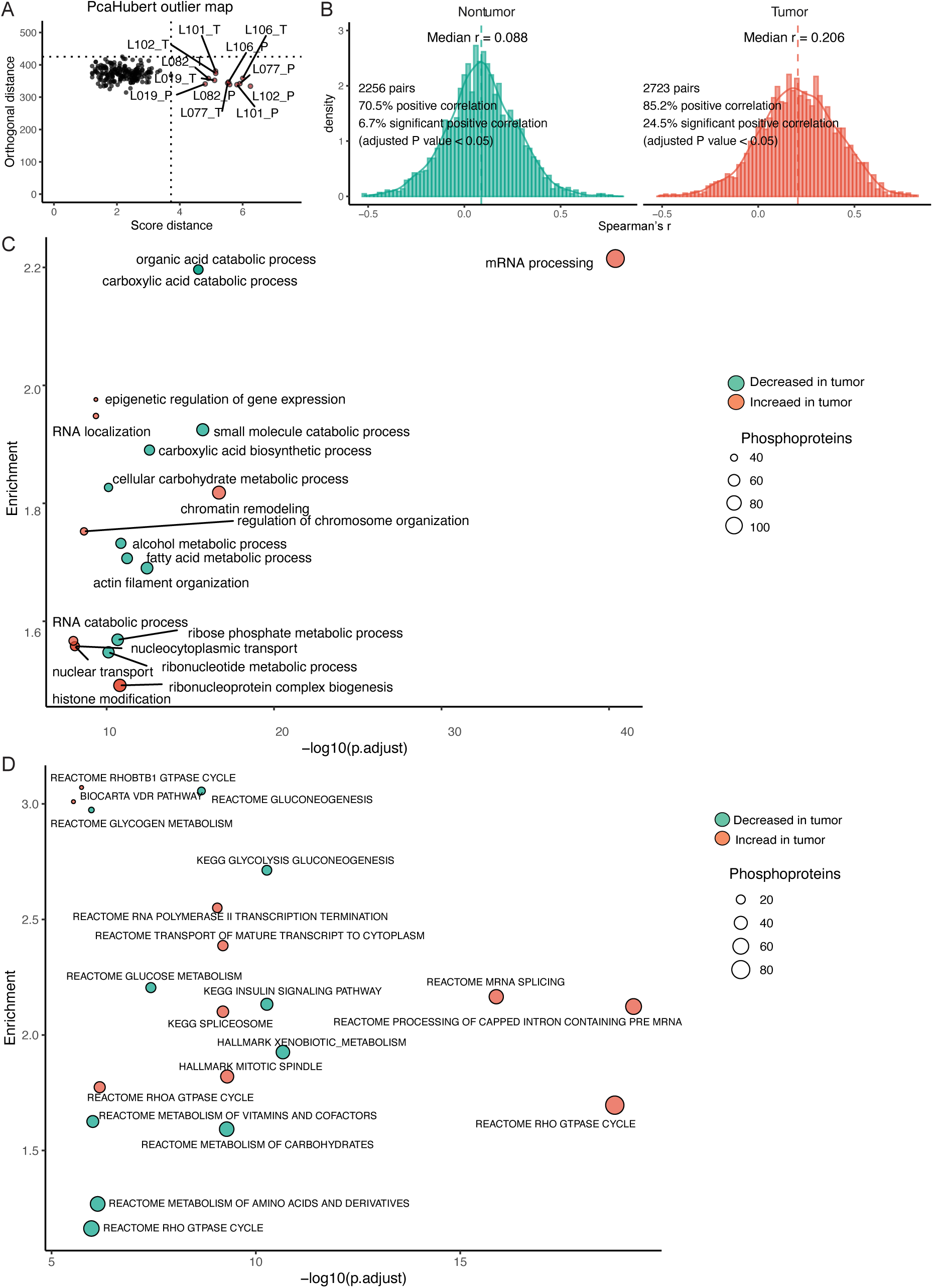
Outlier detection, phosphoproteome and proteome relationship, and dysregulated Gene Ontology terms and pathways in early stage HCC. (A) RobPCA identified 12 outliers (6 paired samples) in the 103 paired non-tumor and tumor samples in Jiang et al.’s cohort. (B) Correlation between protein expression levels and collapsed protein phosphorylation levels in the non-tumor and tumor tissues. (C) The top 10 biological processes of Gene Ontology, (D) pathways and hallmarks with increased and decreased protein phosphorylation levels in tumor compared with non-tumor tissues (Benjamin-Hochberg *adj.p*-value *<*0.05).

**Figure S2.**
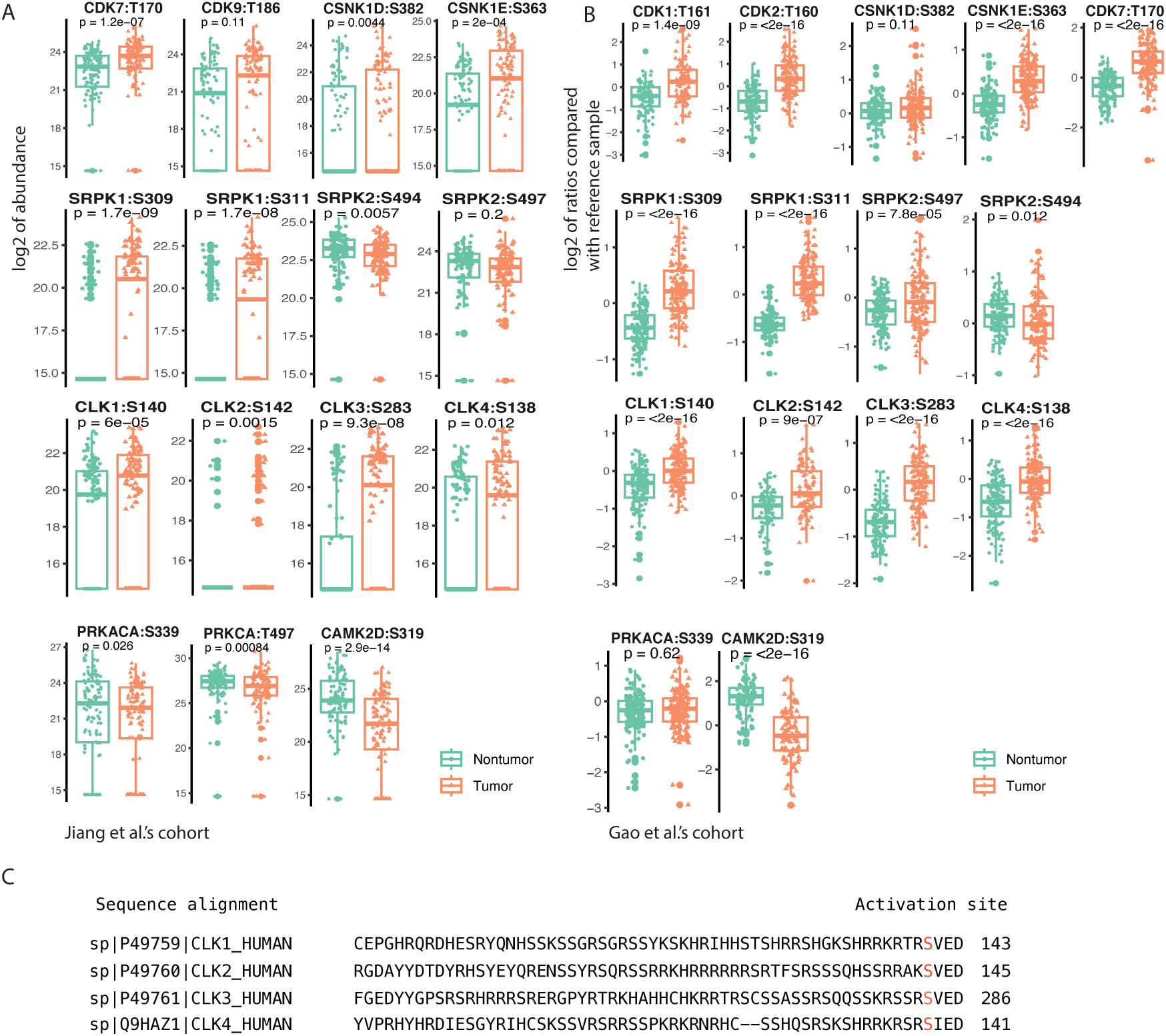
Regulated kinase activities in HCC. (A) Kinase observed with increased and decreased phosphorylation on activation sites in tumor tissues compared to the non-tumor tissues in Jiang et al.’s cohort and (B) Gao et al.’s cohort. The gene name of kinases and the amino acid sequence on their activation sites was labeled in each boxplot. (C) Sequence aligment of CLK1-4 kinases showed the conserved activation sites, which are highlighted in red. Statistical information: the box plot definition is as in Figure 3, *n*=97 for both the non-tumor and tumor samples in panels A and *n*=159 for both the nontumor and tumor samples in panels B.

**Figure S3.**
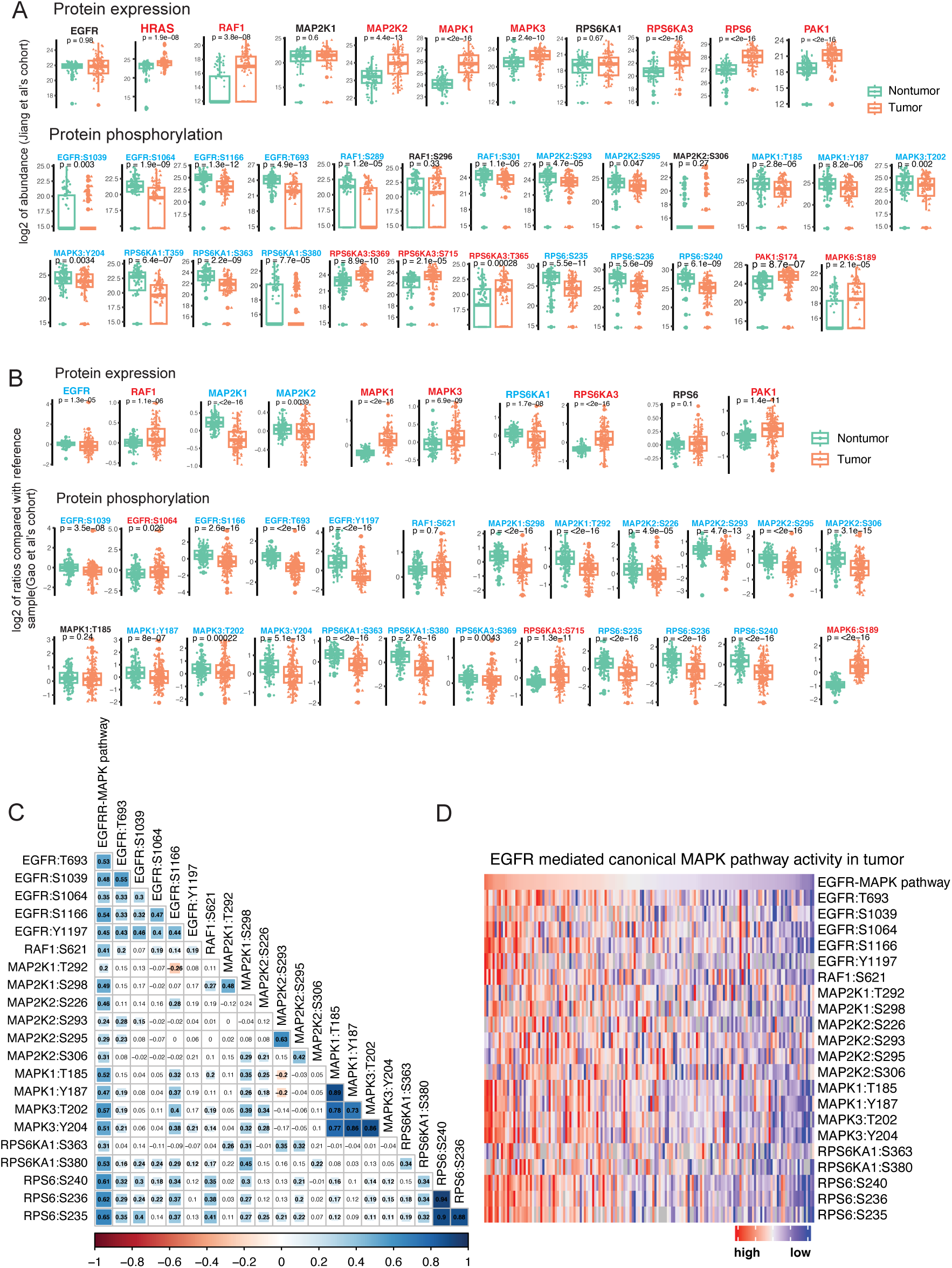
Phosphorylation of the EGFR-mediated canonical ERK/MAPK and atypical MAPK signaling cascades in Jiang et al.’s cohort and Gao et al.’s cohort. (A) The protein expression and phosphorylation levels of key signaling molecules in EGFR-mediated canonical ERK/MAPK and atypical MAPK signaling in HCC and non-tumor tissues in Jiang et al.’s cohort and (B) Gao et al.’s cohort are shown. The expression of proteins and functional phosphosites in non-tumor and tumor tissues are shown in boxplots. Proteins/phosphosites highlighted in red indicate significantly upregulated levels in HCC, while those in blue indicate significantly downregulted levels in HCC. Those in black indicate no significant regulation in HCC. (C) The correlation of the phosphorylation levels of activation sites of proteins in EGFR-mediated canonical ERK/MAPK signaling cascades in Gao et al.’s cohort. The median value of their activation sites was taken as a proxy for the EGFR-mediated canonical ERK/MAPK pathway and (D) patients were ranked from high to low by that and the heatmap of their phosphorylation levels is displayed. Statistical information: Box plot definitions are the same as in Figure 3. In panel A, the sample sizes for both non-tumor and tumor groups are n = 101 for protein expression and n = 97 for protein phosphorylation. In panel B, the sample size is n = 159 per group for both protein expression and phosphorylation.

**Figure S4.**
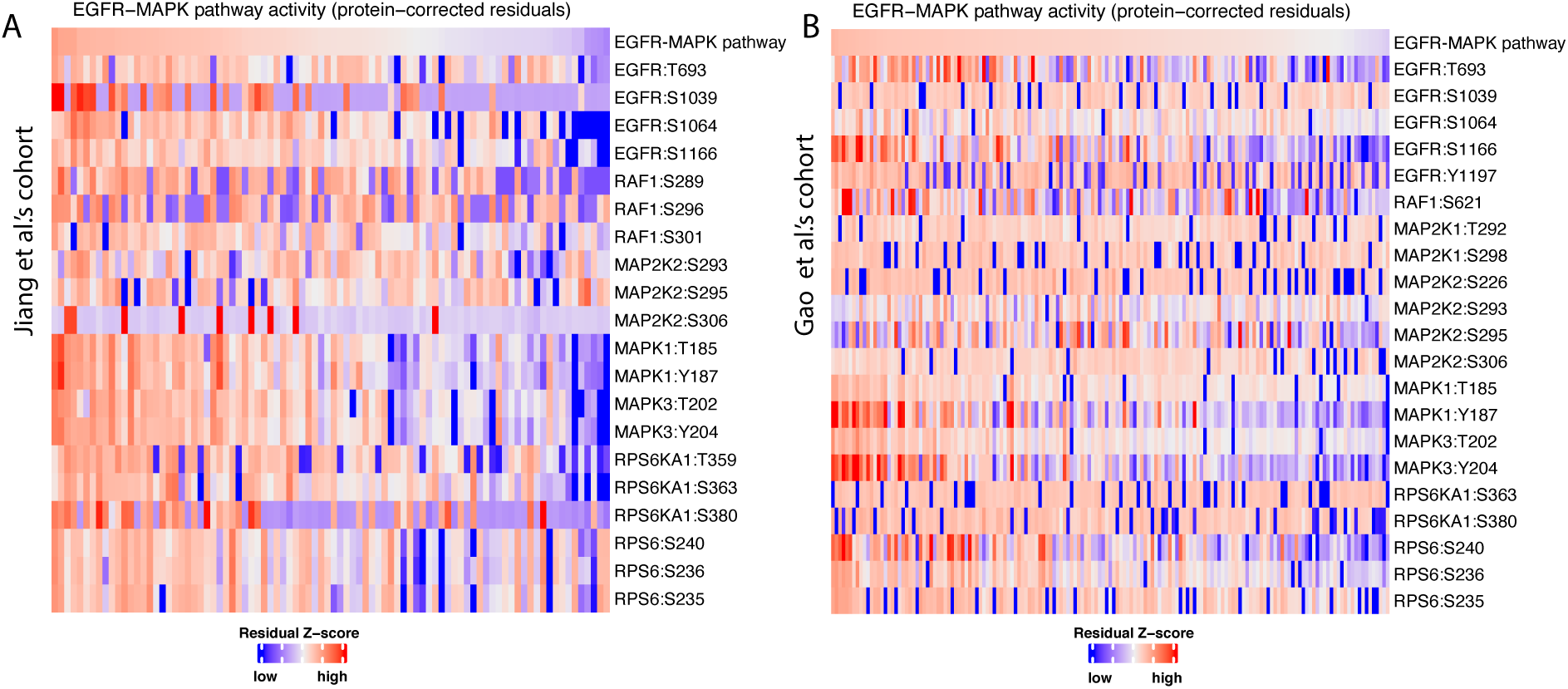
Site-level residuals of EGFR-mediated canonical ERK/MAPK pathway. Linear regression was applied for each phosphosite to regress out its total protein abundance from the phosphosite intensity in (A) Jiang et al.’s cohort and (B) Gao et al.’s cohort, the median residual of their activation sites was taken as a proxy of EGFR-mediated canonical ERK/MAPK pathway. The patients were ranked from high to low by that and the heatmap of their site-level residuals was shown.

**Figure S5.**
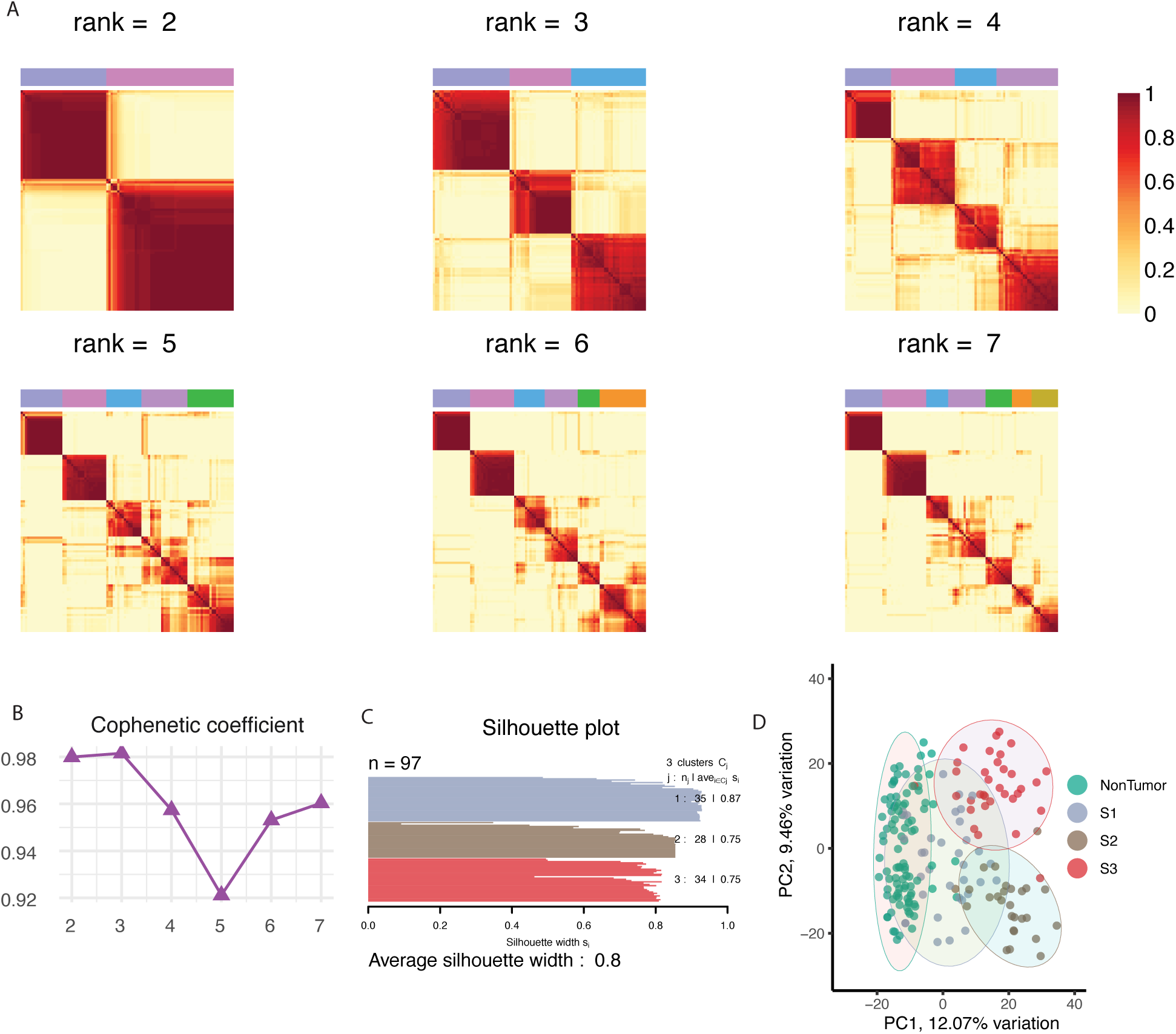
Nonnegative matrix factorization and consensus-clustering identified three phosphoproteomic subtypes in early-stage HCC tissues. (A) NMF and consensus-clustering analysis were applied on 97 samples with rank from 2 to 7 on phosphoproteomic data in Jiang et al.’s cohort [1], and the consensus matrix is shown in the heatmap. (B) Rank = 3 was determined as the final solution since it yieled the largest cophenetic coefficient with (C) average silhouette width of 0.8. (D) Annotation of the three phosphoproteomic subtypes on the scatter plot of principal component analysis of 97 paired samples by the 1,816 phosphoisites, the ellipse presents the 0.95 confidence intervals for each group.

**Figure S6.**
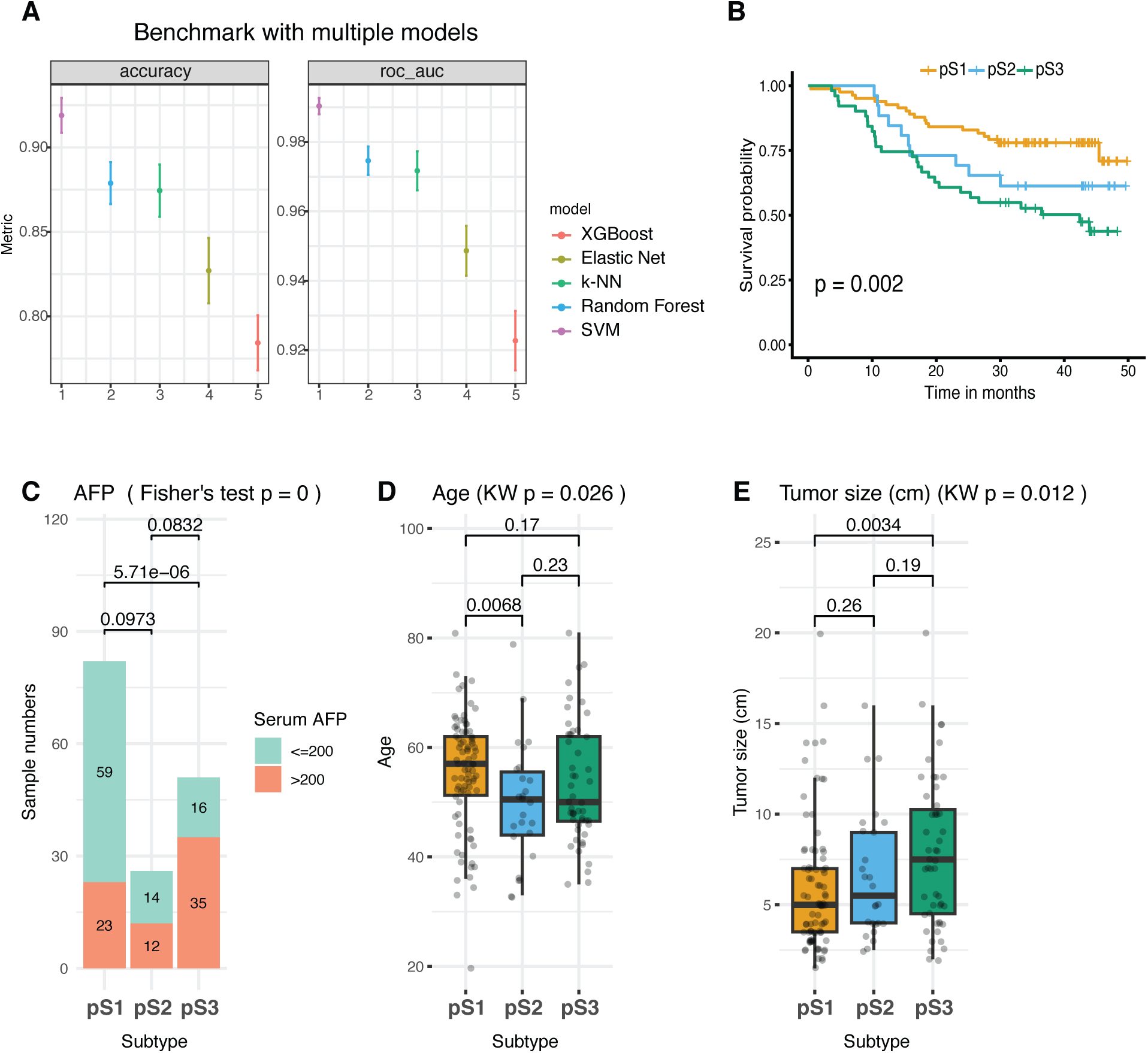
Validation of phosphoproteomic subtypes in Gao et al.’s cohort. (A) Benchmark with multiple machine learning classifiers on Jiang et al.’s cohort, the SVM showed the best performance and was applied on Gao et al’s cohort. (B) Kaplan–Meier curves for overall survival (OS), based on the predicted phosphoproteomic subtypes by SVM model (logrank test). (C) Association of predicted phosphoproteomic subtypes with clinical features, including serum AFP levels, (D) age, and (E) tumor size in Gao et al.’s cohort. Statistical information: the box plot definition is as in Figure 3, The sample sizes of pS1, pS2, pS3 are *n*=82, 26, 51 for panels D and E.

**Figure S7.**
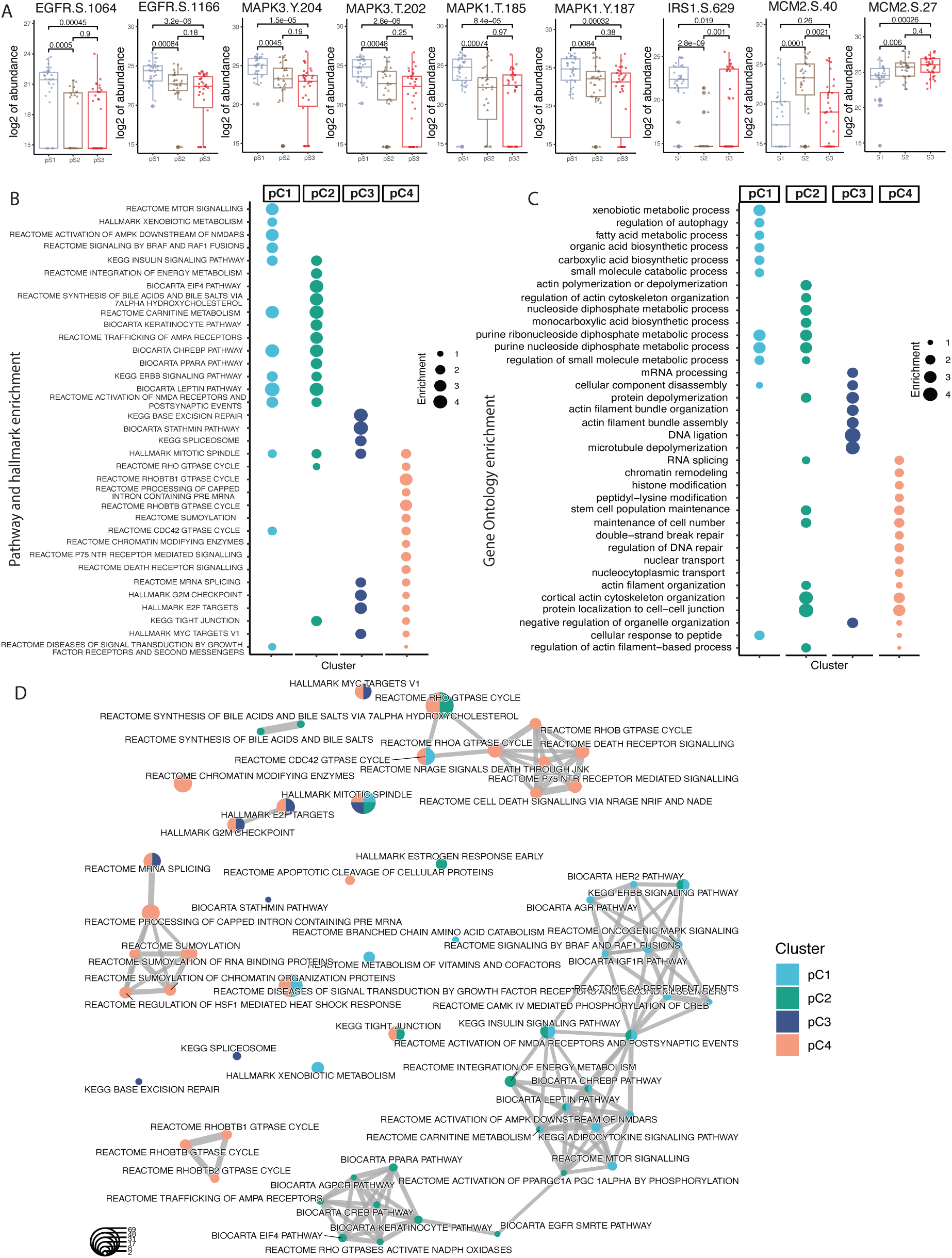
Characterization of representative phosphosite signatures, pathways and hallmarks, biological processes across the three phosphoproteomic subtypes in HCC tissues. (A) Representative phosphosites relating to EGFR-mediated canonical ERK/MAPK signaling, insulin signaling pathway, and DNA replication during cell cycle across the phosphoproteomic subtypes. (B) Clusters of significantly differently expressed phosphosites among the phosphoproteomic subtypes with top 10 pathways and hallmarks, (C) biological processed (Benjamin-Hochberg *adj.p*-value *<*0.05) for each cluster. (D) The emmaplot of top 10 pathways and hallmarks from proteins in four clusters with significantly differently expressed phosphosites across three phosphoproteomic subtypes. Statistical information: Box plot definitions are the same as in Figure 3. The sample sizes in panel A are as in Figure 3D.

**Figure S8.**
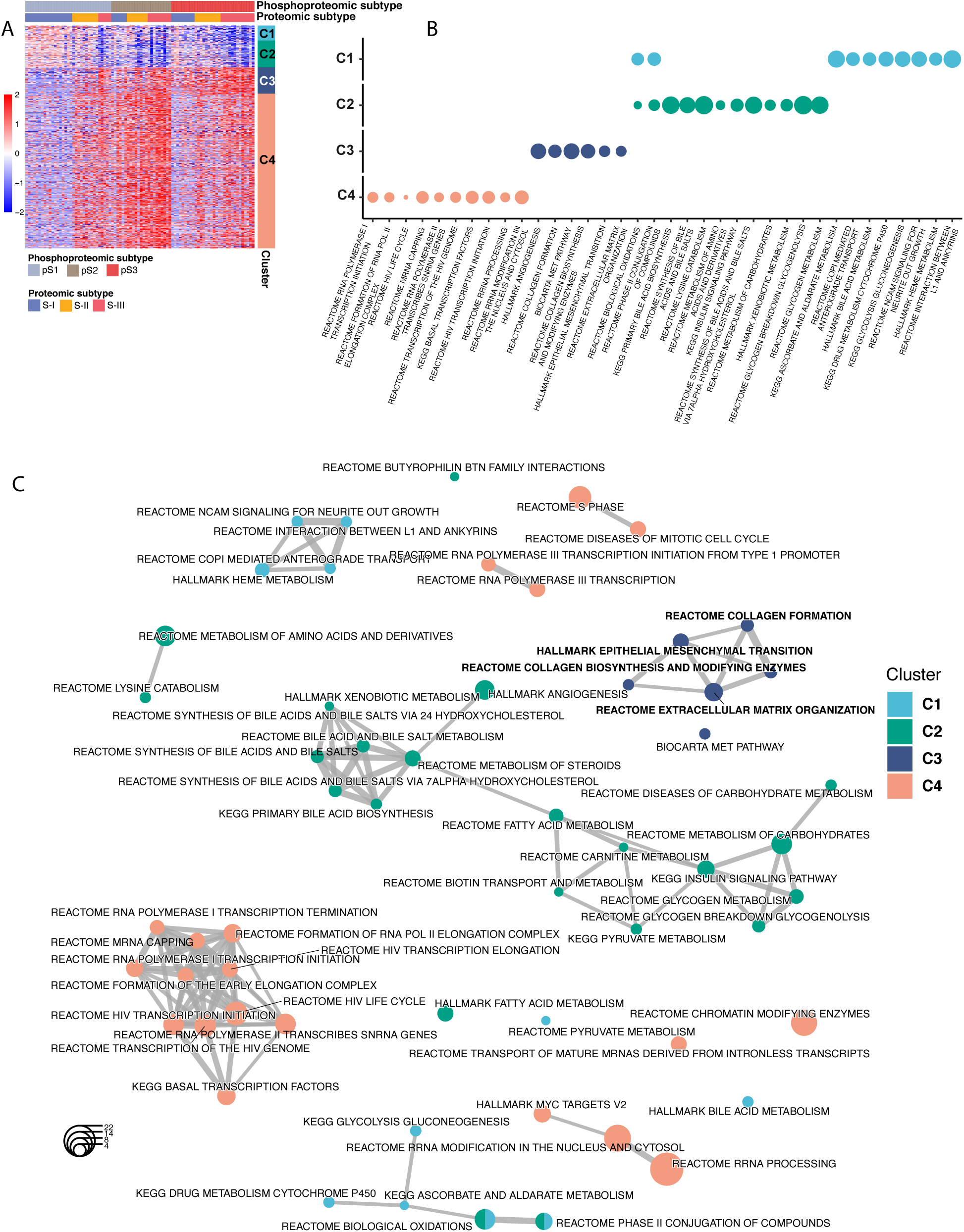
Characterization of pathways and hallmarks of the proteome across the three phosphoproteomic subtypes in HCC tissues. (A) 566 proteins showed significantly different expression levels across the three phosphoproteomic subtypes and were divided into four clusters. The relative abundance of proteins (log2transformed) were shown in the heatmap. (B) Clusters of significantly differently expressed proteins across the phosphoproteomic subtypes with top 10 pathways and hallmarks (Benjamin-Hochberg *adj.p*-value *<*0.05) for each cluster. (C) The emmaplot of top 10 pathways and hallmarks from proteins in four clusters with significantly different expression levels across three phosphoproteomic subtypes. The pathways related to collagen formation, EMT, and ECM were highlighted in bold.

**Figure S9.**
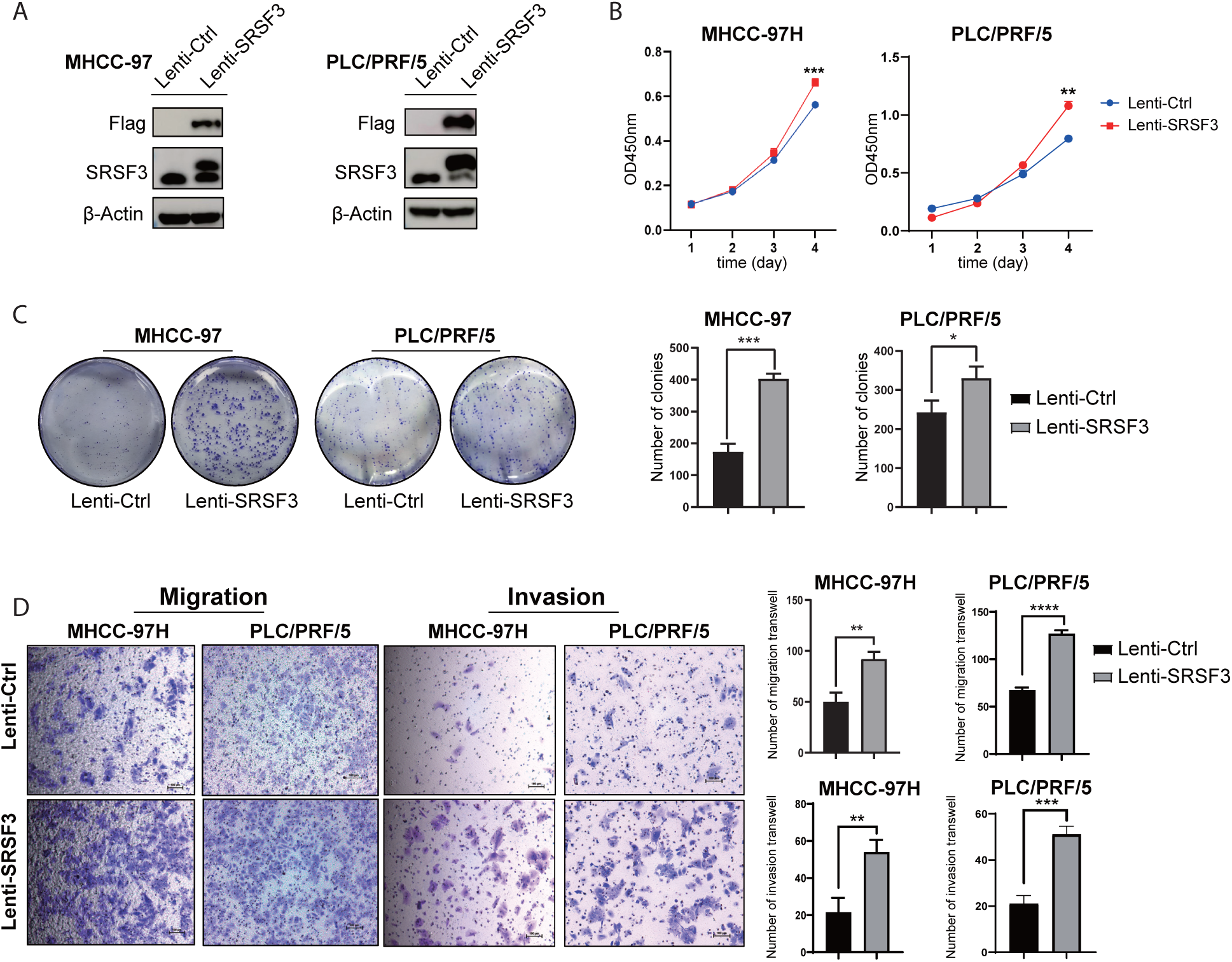
SRSF3 contributes to proliferation, clone formation, migration and invasion of HCC cells. (A) Stable SRSF3 overexpression in MHCC-97H and PLC/PRF/5 cell lines were constructed and significantly enhanced the abilities of (B) proliferation, (C) cell clone formation, (D) migration and invasion of HCC cells. Statistical information: the error bar definition is as in Figure 4. *n*=3 for panels C and D. For all graphs, asterisks show statistical significance by two-sided Wilcoxon rank-sum test, \**p <*0.05; \*\**p <*0.01; \*\*\**p <*0.001.

**Figure S10.**
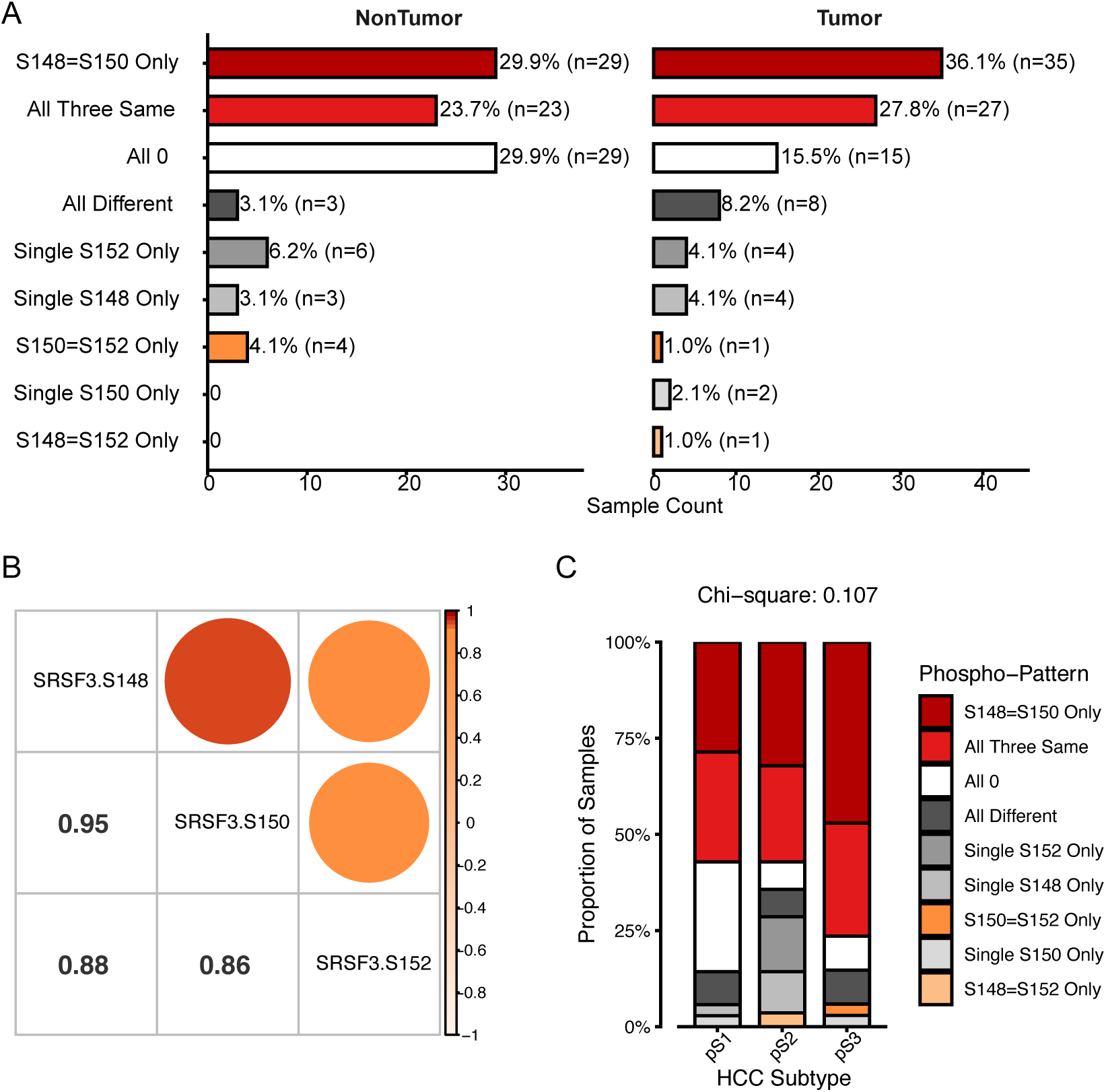
The quantification patterns of phosphorylation of Ser148, Ser150, Ser152 on SRSF3 in the samples. (A) Detailed quantification pattern of the phosphosites on SRSF3 in nontumor and tumor group. ‘S148=S150 Only’: Ser148 and Ser150 were quantified with identical positive intensity value while Ser152 with different value or absence; ‘All Three Same’: Ser148, Ser150, and Ser152 are quantified with identical positive intensity value; ‘All 0’: Ser148, Ser150, and Ser152 are quantified with absence; ‘All Different’: Ser148, Ser150, and Ser152 are quantified with distinct positive intensity value; ‘Single S148/S150/S152 Only’: only one phosphosite was quantified with positive intensity value while the other two with absence. Ser148 and Ser150 showed identical quantification intensity values in 83.5% of tumor cases (‘S148=S150 Only’ + ‘All Three Same’ + ‘All 0’ + ‘Single S152 Only’, *n*=97). (B) Quantification correlation of SRSF3 phosphorylation in tumor. (C) Statistical analysis of SRSF3 phosphorylation patterns among the subtypes.

**Figure S11.**
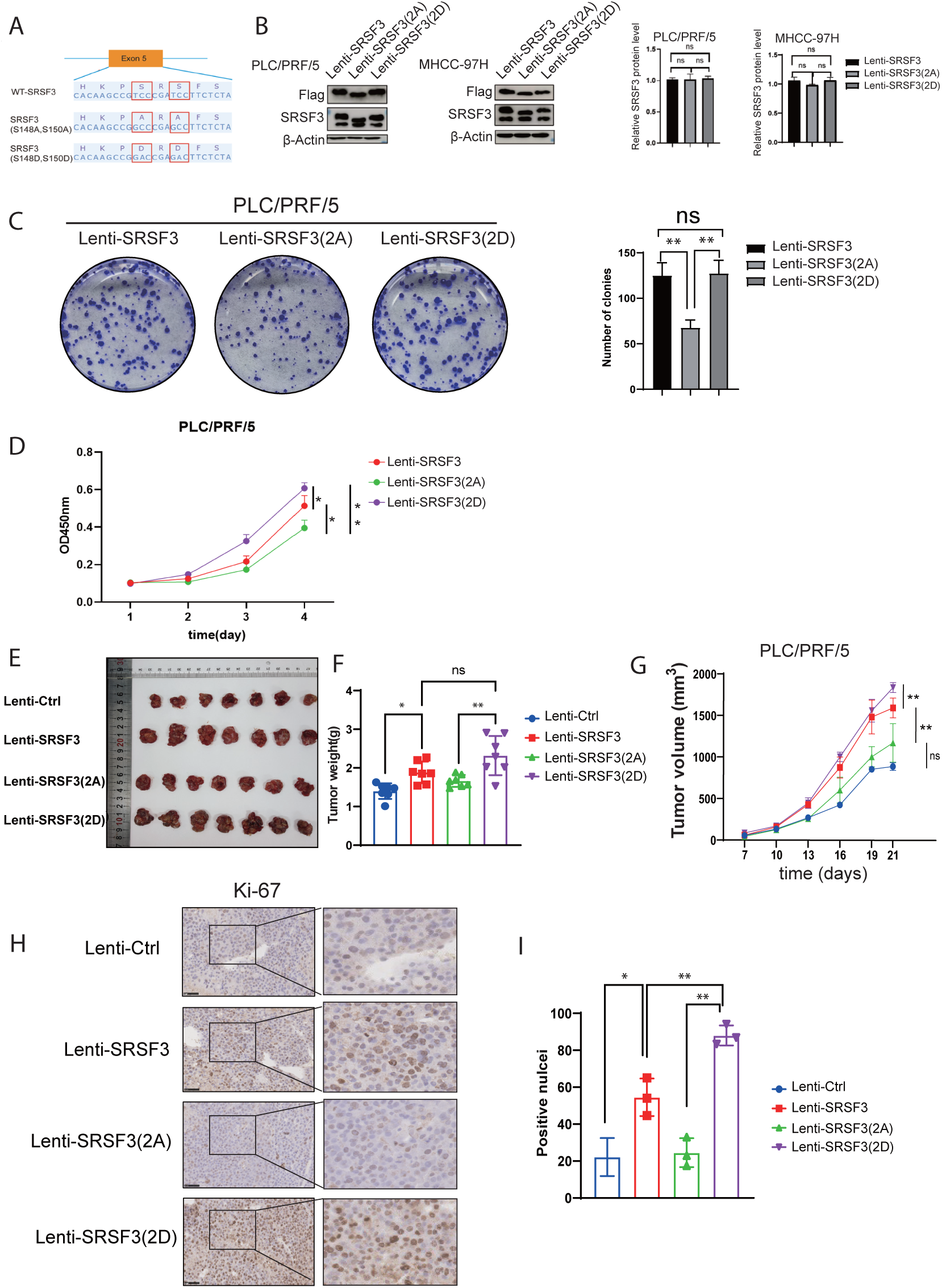
Phosphorylation on Ser148 and Ser150 of SRSF3 promotes proliferation of HCC cells. (A) Stable cell lines were constructed expressing wide-type form, mutant simulating dephosphorylated form (S148A, S150A), mutant simulating phosphorylated form (S148D, S150D) and (B) were confirmed by Western-Blot analysis. (C) Colony formation assays showed that SRSF3 (S148A/S150A) mutant significantly inhibited cell clone formation and (D) proliferation of HCC cells. (E) Images of the gross tumor, (F) the plot of tumor weight and (G) the growth curve from xenograft mouse model in different groups. (H) Immunohistochemistry (IHC) using the Ki-67 antibody highlighted proliferative activity in tumor tissues from the xenograft mouse model in groups with different SRSF3 mutants. (I) Statistical analysis of Ki-67-positive nuclei proportions in groups with different SRSF3 mutants. Statistical information: the error bar definition is as in Figure 4. The sample sizes in panel A are as in Figure 3D, *n*=3 for panels B-D, *n*=7 for panel E-G, *n*=3 for panel I. For all graphs, asterisks show statistical significance by two-sided Wilcoxon rank-sum test , \**p <*0.05; \*\**p <*0.01; \*\*\**p <*0.001.

**Figure S12.**
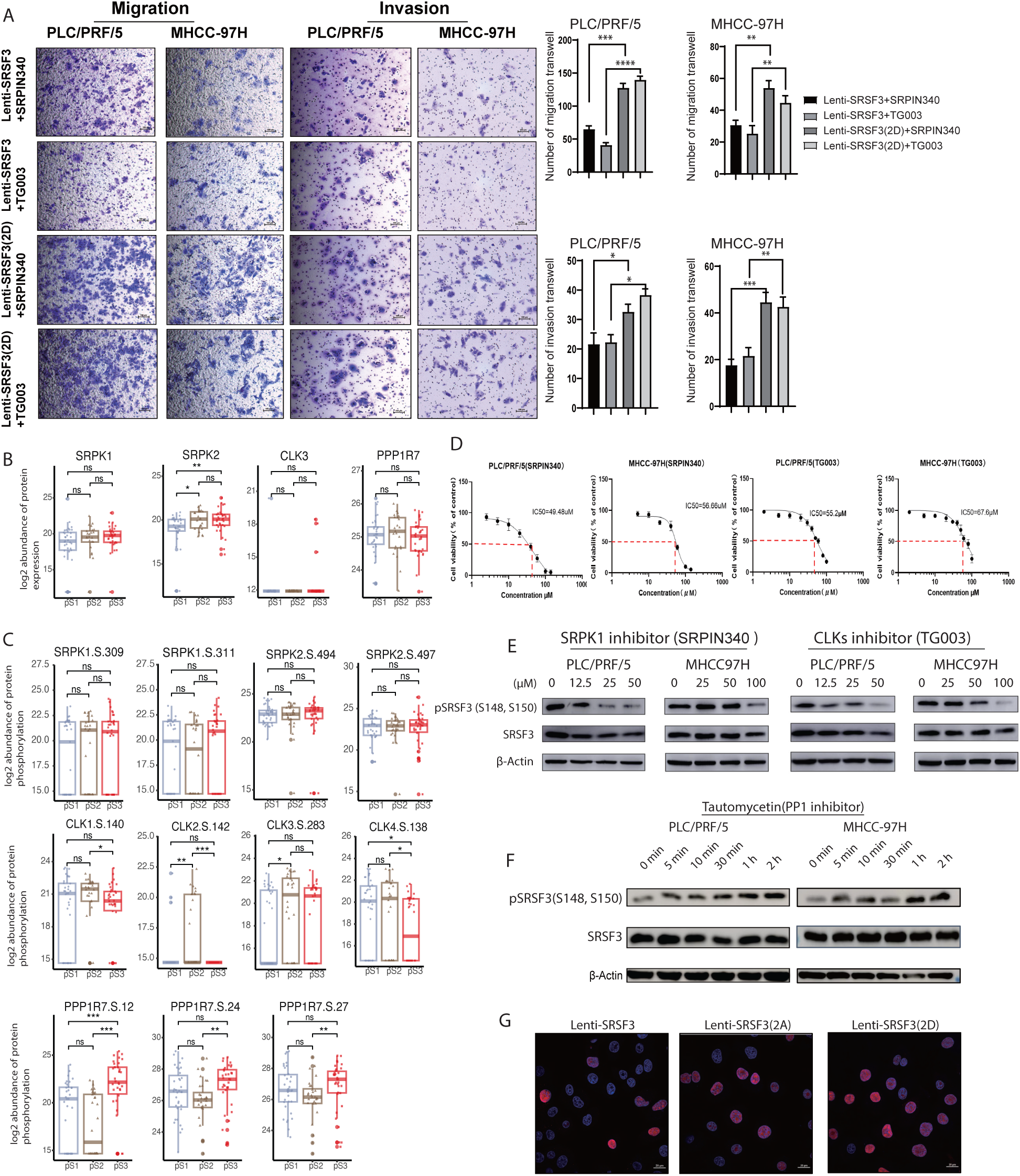
Phosphorylation on Ser148/Ser150 of SRSF3 promotes migration and invasion of HCC cells, regulated by SRPKs/CLKs and PP1. (A) Treatment with SRPIN340 or TG003 was applied to SRSF3-WT overexpressing or phosphorylated mutant SRSF3(S148D/S150D) overexpressing PLC/PRF/5 and MHCC-97H HCC cell lines, respectively. The treatment significantly reduced the migration and invasion abilities of HCC cells in the SRSF3-WT cell lines compared to the phosphorylated mutant. (B) The protein expressions of CLK3, SRPK1, SRPK2 and PPP1R7, and (C) protein phosphorylation level of functional regulating phosphosites on CLKs, SRPKs and PPP1R7 across the three phosphoproteomic subtypes were analyzed. (D) The IC50 of SRPIN340 and TG003 inhibitors in PLC/PRF/5 and MHCC-97H cell lines were determinated. (E) SRPIN340 and TG003 were applied to PLC/PRF/5 cells (0–50 *µ*M for 8 h) and MHCC-97H cells (0–100 *µ*M for 48 h), respectively. Addition of the SRPK1 kinase inhibitor SRPIN340 or the CLKs kinase inhibitor TG003 effectively reduced the abundance of phosphorylation at Ser148 and Ser150 on SRSF3 in these cell models. (F) The addition of tautomycetin (PP1 inhibitor) to PLC/PRF/5 and MHCC97H cell lines significantly increased the abundance of phosphorylation of Ser148 and Ser150 on SRSF3. (G) No significant differences were observed in the subcellular localization of SRSF3 across the SRSF3 mutants. Statistical information: the box plot and error bar definitions are as in Figure 3 and 4. The sample sizes in panel B are *n*=33, 23, 32 for pS1, pS2, pS3 subtype, respectively. The sample sizes in panel C are as in Figure 3D, *n*=3 for panels A and D. For all graphs, asterisks show statistical significance by two-sided Wilcoxon rank-sum test , \**p <*0.05; \*\**p <*0.01; \*\*\**p <*0.001.

### Supplementary tables

**Table S0** Phosphoproteomic data QC status of 103 HCC cases in Jiang et al.’s cohort

**Table S1** GSEA enrichment analysis of pathways and hallmarks for proteins with the high or low correlations of protein expression and phosphorylation level in tumor tissues

**Table S2** Differential expressed phosphorylation sites (DES) analysis on the phosphoproteome in early-stage HCC

**Table S3** Enrichment analysis of biological processes of Gene Ontology for the proteins with differential expressed phosphorylation sites (DES) in early-stage HCC

**Table S4** Enrichment analysis of pathways and hallmarks for the proteins with differential expressed phosphorylation sites (DES) in early-stage HCC

**Table S5** Differential expressed proteins (DEP) analysis on the proteome in early-stage HCC

**Table S6** Enrichment analysis of biological processes of Gene Ontology for the differential expressed proteins (DEP) in early-stage HCC

**Table S7** Enrichment analysis of pathways and hallmarks for the differential expressed proteins (DEP) in early-stage HCC

**Table S8** Pathways and hallmarks enriched from regulated proteins and phosphoproteins in early-stage HCC

**Table S9** Motifs enriched from the upregulated and downregulated phosphosites in early-stage HCC

**Table S10** Kinase activities inffered by KSEA in early-stage HCC

**Table S11** Signature phosphosites for different phosphoproteomic subtypes in early-stage HCC

**Table S12** Pathways and hallmarks enriched from phosphoproteins with signature phosphosites in different cluster in tumor tissues

**Table S13** GO terms enriched from phosphoproteins with signature phosphosites in different cluster in tumor tissues

**Table S14** Pathways and hallmarks enriched from signature proteins in different cluster in tumor tissues

**Table S15** GO terms enriched from signature proteins in different cluster in tumor tissues

**Table S16** Kinase activities inferred by ssKSEA that significantly regulated in the phosphoproteomic subtypes

**Table S17** Site-level residuals after regressing out total protein and pathway activity scores of canonical EGFR-ERK/MAPK signaling in Jiang et al.’s and Gao et al.’s cohorts

## Supporting Methods

### Normalization of phosphoproteomic data and quality assessment

The phosphoproteomic dataset with 27,651 phosphosites (with site localization scores *>*0.75), and the proteomic dataset with 9,254 proteins from 103 tumor and non-tumor paired samples in Jiang et al.’s cohort [1] were processed in the R environment (version 4.2.1). The intensities of phosphosites were *log*_2_ transformed and normalized using the *normalizeQuantiles* function from the R package limma. To detect outliers in the dataset, we applied robust principal component analysis (ROBPCA) to yield a robust estimate of the data [2]. As a result, six paired samples were examined as outliers and were excluded from further analysis (Figure S1 A). The phosphoproteomic dataset with 26,418 phosphosites, and the proteomic dataset with 6,478 proteins from 159 paired tumor and non-tumor samples from Gao et al.’s cohort [3] were used to verify the findings of this study. This dataset was generated using the TMT-multiplexing approach, which inherently minimizes LC-MS/MS batch effects. The ratios of the samples vs. the internal reference were calculated and transformed by *log*_2_, normalized by using median centering to correct for sample loading differences. The phosphosites having more than 50% missing data were excluded, the PCA clearly separated the tumor and normal samples based on the TMT MS data, and no batch effects were observed [3]. We used this rigorously normalized dataset directly.

### Principal component analysis

We performed PCA on 97 paired tumor and non-tumor samples to elucidate the global phosphoproteomic differences between tumor and non-tumor samples. The analysis was restricted to 13,303 phosphosites with quantification coverage of at least 25% in either tumor or nontumor samples individually. The samples were projected onto the first two components, and the 95% confidence intervals for each group were visualized using a colored ellipse.

### Correlations of protein expression and phosphorylation level

The 88 paired tumor and non-tumor samples were analyzed, as they contained both proteomic and phosphoproteomic data. Correlations between protein expressions and phosphorylation levels were calculated separately for tumor and non-tumor tissues. Only proteins or phosphosites quantified in at least 25% of samples were included in the analysis. The protein IDs were mapped to the gene names. Spearman’s correlation coefficients and *p*-values were calculated between phosphosite and protein expression levels. The *p*-values were further adjusted using the Benjamin-Hochberg procedure, and pairs with an *adj.p*-value *<*0.05 were determined as significant. For aggregated phosphoprotein analysis, we first estimated the total phosphorylation level of a given protein by calculating the sum of linear intensities of all phosphosites on the same protein (gene name), followed by a *log*_2_ transformation, and correlations between the aggregated protein phosphorylation levels and the corresponding protein abundance were calculated as described above.

To investigate the pathways and hallmarks associated with high or low correlations between protein expression and phosphorylation levels, gene sets from the R package msigdbr (version 7.5.1) were used. These included 186 Kyoto Encyclopaedia of Genes and Genomes (KEGG) gene sets, 292 Biocarta gene sets, 1615 Reactome gene sets, 196 PID gene sets, 10 NABA gene sets, and 50 cancer hallmark gene sets. Proteins were ranked based on their Spearman’s correlation coefficients (from high to low), and the *GSEA* function in the R package clusterProfiler (version 4.8.3) was applied for the enrichment analysis. The minimum gene set size (minGSSize) was set to 10, and *p−*values were adjusted using the Benjamin-Hochberg procedure, and an *adj.p−*value cut-off of 0.05 was used as the significance threshold.

### Differentially expressed phosphosites (DES) and proteins (DEP) between tumor and non-tumor

The protein expression and phosphorylation values were *log*_2_ *−transformed* and normalized using the *normalizeQuantiles* function in the R package limma. After filtering with a coverage cutoff *≥* 50% in at least one group of tumor or non-tumor tissues, missing values were imputed by the global minimum in the entire dataset. We applied this strategy consistently with the original Jiang et al.’s methodology [1], which is a standard left-censored approach for imputing the missing values in shotgun proteomic datasets [4]. To identify the significantly regulated phosphosites/proteins between early-stage HCC and paired non-tumor samples, a linear model was fitted using *limma*, accounting for within-patient correlations estimated by the *duplicateCorrelation* function. *Moderated t-statistics* were implemented, and *p*-values were adjusted by the Benjamini-Hochberg procedure to control the false discovery rate. Following differential analysis, a total of 3,610 DESs and 3,628 DEPs were observed with an *adj.p*-value *<*0.05 and a *|log*_2_ (fold change)*| >*1.

### Motif enrichment and kinase-substrate enrichment analysis (KSEA)

The significantly elevated and decreased DESs were used for motif enrichment and kinase activity prediction, respectively. For motif enrichment, the Motif-X algorithm [5] was used with a *p*-value cutoff of 1e-5, minimum sequence length (min.seq) =20 for the amino acids ‘S’, ‘T’, and ‘Y’, and the sequences of all phosphosites observed in the dataset were used as the background sequences, respectively. For KSEA, the enzsub database was downloaded from Omnipath[6]. The DESs together with the phosphosites not significantly regulated (n = 7,575) were used and ranked by -log10(p-value) * sign(coef(x)), and the *GSEA* function with minGSSize=5, nPerm=10,000 was applied to infer the kinase activities. A *q* -value less than 0.25 was considered the cut-off. Only kinases detected with regulated phosphosites in HCC were considered as kinases with elevated or decreased activities in HCC.

### Biological process and pathway alteration on protein expression and phosphorylation level

The *enrichGO* function in the R package clusterProfiler was used for Gene Ontology (GO) enrichment analysis. For pathway and hallmark enrichment analysis, the KEGG, Biocarta, Reactome, PID, NABA gene sets, and cancer hallmark gene sets in the migsdb database in the R package msigdbr combined with the function *enricher* in the R package clusterProfiler were applied. For GO, pathway and hallmark enrichment results, *p*-values were adjusted using the Benjamin-Hochberg procedure, with a cut-off of 0.05. The similarity in the pathway and hallmarks enrichment was calculated using the Jaccard correlation coefficient (JC) method, and the enrichment map was visualized using the function *emapplot* in clusterProfiler.

### UpSet plots of pathways and hallmarks

To visualize the overlap between pathways and hallmarks, UpSet plots were created using the R package UpSetR.

### Identification of phosphoproteomic subtypes

Nonnegative matrix factorization (NMF) is widely used for dimension reduction and has demonstrated its ability to extract biologically meaningful information from high-dimensional data[1, 7]. To uncover the subtypes in early-stage HCC, we first filtered out phosphosites quantified in less than 20% of the tumor samples. The median absolute deviation (MAD) was calculated for each phosphosite across samples. Subsequently, the bottom 20% of phosphosites with low MAD were removed, leaving 11,083 phosphosites for NMF factorization. The *nsNMF* algorithm in the R package NMF (v.0.25) was used for subtyping phosphoproteomic data. We evaluated the rank from 2 to 7, using 100 iterations for the rank survey and 500 iterations for unsupervised consensus clustering. Based on cophenetic scores and consensus membership heatmaps, we selected rank=3 as the optimal subtype solution for the phosphoproteomic data (Figure S5).

### Associations between phosphoproteomic subtypes and clinical characteristics

For the associations between phosphoproteomic subtypes and clinical characteristics, we first applied the Kruskal-Wallis (KW) test (for continuous data) or global Fisher’s exact test (for categorical data) to assess global variance across the subtypes. Only when the global test is significant (*p <*0.05), the Wilcoxon rank-sum tests or Fisher’s exact test were evaluated between subtypes. Log-rank test and Kaplan–Meier survival curves were applied to compare the overall survival among the phosphoproteomic subtypes, and a significance level of 0.05 was used as the cut-off threshold.

### Signature phosphosites for phosphoproteomic subtypes and pathways/hallmarks, biological processes, and kinase activities across the subtypes

To identify the signature phosphosites of phosphoproteomic subtypes, we compared the abundance of phosphosites in each phosphoproteomic subtype with that in other subtypes and adjacent non-tumor groups. Phosphosite coverage with a quantification cut-off of 50% in at least one phosphoproteomic subtype or adjacent non-tumor groups was required, and the missing values were imputed with the global minimum value. A linear model was fitted in *limma* by considering the within-patient correlation estimated by the *duplicateCorrelation* function, *moderated t-statistics* were implemented, and *p*-values were adjusted by the Benjamin-Hochberg procedure. To identify signature phosphosites/proteins of the phosphoproteomic subtypes, the following criteria were required: all *adj.p*-values *<*0.05 and at least a two-fold change (lfc=1) when compared to other subtypes and adjacent non-tumor samples. A total of 1,816 phosphosites and 566 proteins were identified as the signature phosphosites/proteins, which were further divided into four clusters (pC1-pC4 for phosphosites and C1-C4 for proteins): the pC1/C1 cluster comprises those with the highest abundance in the pS1 subtype, the pC2/C2 cluster exhibits the lowest abundance in the pS2 subtype, the pC3/C3 cluster shows the highest abundance in the pS3 subtype, and the pC4/C4 cluster shows the highest abundance in the pS2 subtype.

For the enrichment of biological processes, pathways/hallmarks of the clusters, the protein IDs (of phosphosites) in each cluster were used for analysis. The *enrichGO* function in the R package clusterProfiler was used for biological process enrichment analysis. The KEGG, Biocarta, Reactome, PID, NABA gene sets, and cancer hallmark gene sets in migsdb database in the R package msigdbr combined with the function *enricher* in the R package clusterProfiler were used for the pathway and hallmark enrichment analysis. For biological processes, pathway and hallmark enrichment results, the *p*-values were adjusted using the Benjamin-Hochberg procedure and with a cut-off of 0.05. The similarity in the pathways/hallmarks enrichment was calculated using the Jaccard correlation coefficient (JC) method, and the enrichment map was visualized using function *emapplot* in clusterProfiler. For the dotplot of the enrichment results, the enrichment factor was calculated as the ratio of two proportions, that is, Enrichment=log2(ProteinRatio(k/n)/BgRatio(M/N)), where ProteinRatio(k/n) represents the ratio of input proteins that are annotated in a term, and BgRatio (M/N) represents the ratio of all proteins that are annotated in this term.

For kinase activities in the subtypes, the enzsub database from Omnipath[6] was used as a resource for kinase-substrate relation sets to infer kinase activities. We applied single-sample kinase-substrate relation set enrichment analysis (ssKSEA) to infer the kinase activities in each sample, including tumor and non-tumor tissues. This analysis utilized the *gsva* algorithm with the *ssgsea* method from the R package GSVA [8]. The kinase scores reported by the *gsva* algorithm were used as a proxy for kinase activities, and only those observed in the phosphoproteomic data were retained to infer kinase activities in the phosphoproteomic subtypes. By applying the same procedure with signature phosphosites in the subtypes, as described above, and including the kinases that were elevated or decreased in the tumor compared to non-tumor tissues across all three phosphoproteomic subtypes, we identified 57 kinases whose activities were significantly regulated.

### Cross-cohort validation of phosphoproteomic subtypes

We utilized the Gao et al.’s cohort to validate the robustness of the phosphoproteomic subtypes. Given the differences in phosphopeptide enrichment (TiO_2_ *vs.* IMAC) and quantification (DDA label-free *vs.* TMT-multiplexing DDA) between the Jiang et al’s and Gao et al.’s cohorts, we implemented a robust supervised machine learning validation workflow. Of the 1,816 phosphosites discovery signatures that were significantly regulated among the phosphoproteomic subtypes, 1,334 phosphosites were successfully quantified in Gao et al.’s dataset. To ensure a balanced representation of the four molecular clusters (pC1–pC4) of phosphosite signatures, we determined the feature size (*n* = 128) based on the smallest cluster (pC3) in the Gao et al.’s cohort. We then selected the top 128 phosphosites of each cluster, ranked by their fold-change coefficients, which resulted in a panel of 512 phosphosites. The quantification value for this final panel was extracted from both datasets, and z-score scaling across the tumor samples within each cohort was applied to mitigate batch effects. We then benchmarked five machine learning classifiers (SVM, Random Forest, XGBoost, Elastic Net, and k-NN) using 10 repeats of 5-fold cross-validation on the discovery cohort. The model with the highest internal accuracy and ROC-AUC was further applied to Gao et al.’s cohort and stratified the patients.

### Lentivirus-infected cells

When the cells reached 60%-70%, the medium was changed to serum-free DMEM or MEM, and lentivirus and polybrene were added to each well. Then the cells were incubated. After 24 hours of infection, the medium was replaced with a complete medium. For Huh-7 and MHCC-97H cells knockdown, puromycin (working concentration 2 *µ*g/mL, Shanghai Taosen Biochemical Technology) was used for screening over 72-96 hours. For MHCC-97H and PLC/PRF/5 cells overexpressing SRSF3 and phosphomutants, Geneticin (G418) (working concentration 800 *µ*g/mL, Beyotime Biotechnology) was used for screening for at least one week.

### Design and synthesis of SRSF3 protein phosphomutants and interference sequences

The nucleotide sequence of SRSF3 was obtained from the NCBI database. Lentivirus vectors LV-EFS-Kozak-Flag-SRSF3 (Ser148, Ser150), LV-EFS-Kozak-Flag-SRSF3 (S148A, S150A), and LV-EFS-Kozak-Flag-SRSF3 (S148D, S150D), as well as the control group LV-EFSKozak-Flag-NC, were constructed by Sigma (Suzhou) Biotechnology Co., Ltd. S148A (TCC to GCC), S150A (TCC to GCC), S148D (TCC to GAC), and S150D (TCC to GAC) mutations were introduced. The knockdown vectors, LV2-SRSF3-shRNA-PURO and control group LV2-PURO-NC, were constructed by Suzhou Gima Company. The interfering sequence of SRSF3 was 5’-CAGGTCCCTTTCTAGAGATAG-3’.

### Extraction of total cellular protein and western blot

An appropriate volume of 4% SDS lysis buffer was added to the cell pellet, which was incubated on ice for 20 minutes. Ultrasonic cell disruption was performed, and after centrifugation at 12,000 rpm, 4 °C for 20 minutes, the supernatant was collected for determination of protein concentration. For western blotting, the samples were denatured at 95 °C for 5 min, resolved on a 12% SDS-PAGE gel, and transferred to a nitrocellulose membrane. The membrane was blocked with 5% skim milk was used, followed by incubation with the following antibodies: anti-beta-actin (Beijing Zhongshan Jinqiao Biotechnology Co., Ltd.), anti-SRSF3 (Invitrogen, USA), anti-Ki67 (Abcam Company), anti-SRSF3 (S148, S150) phosphorylated antibody (prepared by Wuhan Dion Biotechnology Co., Ltd.), FlAG antibody (Proteintech Company), E-cadherin antibody (CST Company USA), anti-N-cadherin, anti-Vimentin (Proteintech Company), and goat anti-mouse/anti-rabbit secondary antibody ( Beijing Kangwei Century Co., Ltd.).

### Colony formation assay

Cells were seeded in 6-well plates at a density of 800-1,000 cells per well, and the cell culture plates were cultured in triplicate at 37 °C for 8 to 10 days. The cells were then fixed in 4% paraformaldehyde for 15 min and stained with crystal violet for 15 min at room temperature. After washing with PBS, colonies were counted to determine if there were any significant differences.

### Cell proliferation

A CCK-8 assay was used to detect cell proliferation. The cells were plated in a 96-well plate at a density of 6 *×* 10^3^ cells per well (100 *µ*L total volume). After culturing in an incubator for 5 hours to allow adhesion, the 96-well plate marked for 0h was taken out, the medium was discarded, and prepared 10% CCK-8 medium was added. The plate was incubated for one hour in the cell culture box. A microplate reader (Biotek USA) was used to measure the absorbance at OD 450 nm. Absorbance values were recorded at 24h, 48h, 72h, and 96h, and the data were saved.

### Cell migration and invasion

The cells were starved overnight. Subsequently, 600 *µ*L of complete medium was added to the lower compartment of a 24-well plate, and approximately 8*×*10^4^ to 10*×*10^4^ cells/100 *µ*L of pre-calculated cell suspension was added to the upper compartment of each well. Three replicates for each control and experimental group. The cells were incubated in a cell culture incubator for 24 or 48 hours. After removal from the chamber, cells were fixed, stained, and photographed.

### Confocal laser scanning microscopy (CLSM)

Cells were fixed with 4% paraformaldehyde for 15 minutes and washed with PBS. Blocking was performed using 10% normal goat serum for 40 minutes. Incubation with the primary antibody diluted in 5% goat serum was performed overnight at 4°C. The next day, the cells were incubated with the secondary antibody, stained for nuclei, mounted, and imaged.

### In vivo xenograft studies

Male NOD/SCID mice (Beijing Vital River Laboratory Animal Technology Co., Ltd., Beijing, China) aged 5 weeks were housed in a strictly specified pathogen-free (SPF) animal facility. Stable NC, SRSF3-WT, SRSF3-S148A/S150A, and SRSF3-S148D/S150D PLC/PRF/5 cells (5 10^6^) were injected subcutaneously into the right flank of each NOD/SCID mouse. Tumor dimensions were measured every four days using a caliper after tumor formation. The tumor volume was calculated using the formula: 0.5 length (L) width (W)^2^. The mice were euthanized at the end of the trials, and the solid tumors were dissected and weighed.

### Data collection and statistical analysis

The data provided in this study were statistically analyzed in the R environment or using GraphPad Prism 9 software and presented as mean *±* standard deviation (mean *±* SD). Two-sample comparisons were performed using the two-sided Wilcoxon rank-sum test, with *p*-values *<*0.05 considered significant. \**p <*0.05; \*\**p <*0.01; \*\*\**p <*0.001; \*\*\*\**p <*0.0001.

### Dataset acquisition

The proteomic and phosphoproteomic datasets from Jiang et al. [1] and Gao et al. [3] were analyzed in this study, which can be obtained from the iProX database (www.iprox.org, accession number IPX0000937000) and NODE database (https://www.biosino.org/node) by pasting the accession (OEP000321), or through the Bitbucket repository.

